# Rules for designing protein fold switches and their implications for the folding code

**DOI:** 10.1101/2021.05.18.444643

**Authors:** Yingwei Chen, Yanan He, Biao Ruan, Eun Jung Choi, Yihong Chen, Dana Motabar, Tsega Solomon, Richard Simmerman, Thomas Kauffman, D. Travis Gallagher, John Orban, Philip N. Bryan

## Abstract

We have engineered switches between the three most common small folds, 3α, 4β+α, and α/β−plait, referred to here as A, B, and S, respectively. Mutations were introduced into the natural S protein until sequences were created that have a stable S-fold in their longer (∼90 amino acid) form and have an alternative fold (either A or B) in their shorter (56 amino acid) form. Five sequence pairs were designed and key structures were determined using NMR spectroscopy. Each protein pair is 100% identical in the 56 amino acid region of overlap. Several rules for engineering switches emerged. First, designing one sequence with good native state interactions in two folds requires care but is feasible. Once this condition is met, fold populations are determined by the stability of the embedded A- or B-fold relative to the S-fold and the conformational propensities of the ends that are generated in the switch to the embedded fold. If the stabilities of the embedded fold and the longer fold are similar, conformation is highly sensitive to mutation so that even a single amino acid substitution can radically shift the population to the alternative fold. The results provide insight into why dimorphic sequences can be engineered and sometimes exist in nature, while most natural protein sequences populate single folds. Proteins may evolve toward unique folds because dimorphic sequences generate interactions that destabilize and can produce aberrant functions. Thus, two-state behavior may result from nature’s negative design rather than being an inherent property of the folding code.

**Significance Statement:** We establish general rules for designing protein fold switches by engineering dimorphic sequences that link the three most common small folds. The fact that switches can be engineered in arbitrary and common protein folds, sheds light on several important questions: 1) What is the generality of fold switching? 2) What types of folds are amenable to switching? 3) What properties are shared by sequences that can fold into two completely different structures? This work has implications for understanding how amino acid sequence encodes structure, how proteins evolve, how mutation is related to disease, and how function is annotated to sequences of unknown structure.

**Classification:** Biological Sciences: Biochemistry; Physical Sciences: Biophysics and Computational Biology

## Introduction

Fold switching occurs when one amino acid sequence has a propensity for two completely different, but well-ordered, conformations. Many examples of both natural and engineered fold switching demonstrate that proteins can have a stable native fold while simultaneously hiding latent propensities for alternative states with new functions (1-7). This fact has many implications for understanding how amino acid sequence encodes structure, how proteins evolve, how mutation is related to disease, and how function is annotated to sequences of unknown structure. Even as structure prediction has improved, however, detection of latent propensities in a given sequence and prediction of fold switches is usually problematic.

In this paper, we seek to establish some general rules for designing fold switches by engineering switches between three common folds. Our previous studies examined switches between the 3α (G_A_) and the 4β+α (G_B_) domains of Protein G and demonstrated that protein structure can be encoded by a small number of essential residues, and a very limited subset of intra-protein interactions can tip the balance from one fold and function to another (8, 9). Here we determine that both G_A_ and G_B_ can switch into a third fold (α/β−plait), thus connecting three folds in mutational pathways that avoid unfolded states. The premise of this paper is that if switches can be engineered in arbitrary and common protein folds, it will shed light on several important questions: 1) What is the generality of fold switching? 2) What types of folds are amenable to switching? 3) What properties are shared by sequences that can fold into two completely different structures?

The proteins used in this study have no significant homology, represent the three most common fold types (10), and are models for studying protein folding and stability (**Fig. S1**). All are small and amenable to NMR studies. They did not pass any initial test of likely switching. By studying small proteins that are widely used in experimental and computational folding studies, experimental results connect a large body of knowledge e.g. (11-24). Streptococcal Protein G contains two types of domains that bind to serum proteins in blood: the G_A_ domain binds to human serum albumin (HSA) (25, 26) and the G_B_ domain binds to the constant (Fc) region of IgG (27, 28). The ribosomal protein S6 from *Thermus thermophilus* is a well-studied member of the α/β−plait family (29-33). For simplicity, the S6 fold is referred to as an S-fold. When a switch to either the G_A_ or G_B_ fold is discussed, both are referred to as a G-fold. The specific G_A_ fold is referred to as an A-fold and the specific G_B_ fold is referred to as a B-fold.

The basic challenge in designing fold switches is, given two arbitrary folds, how do you identify one sequence that contains the essential folding information for both folds? Theoretically, a simple approach is an exhaustive computational search to find one sequence that has mutually compatible native interactions in two conformations. A more practical design process requires a method for aligning the sequences for the two folds such that essential folding information for both folds can be introduced by mutation in a way that is mutually compatible. For example, automated alignment can thread the shorter sequence through the larger structure and calculate the energy of hypothetical structures in every register (34). This approach is not dependable for designing switches, however, because some alignments with high energies can be radically improved with a few strategic mutations. To maximize possibilities of a parsimonious switch, we aligned the sequences in all registers and evaluated what would have to change to accommodate both folds. The process was as follows:

i. Thread a G-sequence through the S-fold in each possible alignment.
ii. Identify alignments that minimize the number of catastrophic interactions.
iii. Determine tolerable mutations in the G-sequence that might resolve the catastrophic clashes in the S-fold. Redesign clusters of amino acids to resolve clashes. Use the Rosetta-Relax protocol to adjust the peptide backbone and evaluate the energy of the design (35).
iv. Optimize protein stability in the S-fold by computationally mutating amino acids at non-overlapping positions. Repeat minimization and evaluation with Rosetta-Relax. To minimize uncertainties involved in computational design, conserve original amino acids whenever possible.

Previously we created sequences that populate both A- and B-folds by threading the A-sequence through the B-fold, finding a promising alignment, and then using phage-display selection to reconcile one sequence to both folds (8, 36, 37). Here the approach is conceptually similar, except that we use Rosetta as a computational design tool to identify compatible mutations rather than phage display. There is no reason to assume that this method is optimal. We are merely applying a practicable scheme for engineering dimorphic sequences and then evaluating the outcome using structure determination by NMR and thermal denaturation.

Five dimorphic sequence pairs (10 proteins) were designed and purified (**Fig. 1**). In each of these designs, the protein pairs are 100% identical in a 56 amino acid region of overlap. Analysis of thermal denaturation showed eight of the 10 to be stable, well-populated structures. Seven structures were determined using NMR spectroscopy and compared to the designs. Two of the switches (one S-to A-switch and one S-to B-switch) achieved the goal of populating the S-fold in the longer form and the A- or B-fold in the shorter form. The other cases were equally informative, however. Here, we describe the folding energetics and structures of these 10 dimorphic proteins and present a set of basic principles for designing fold switches that emerged from this analysis.

**Figure 1:**
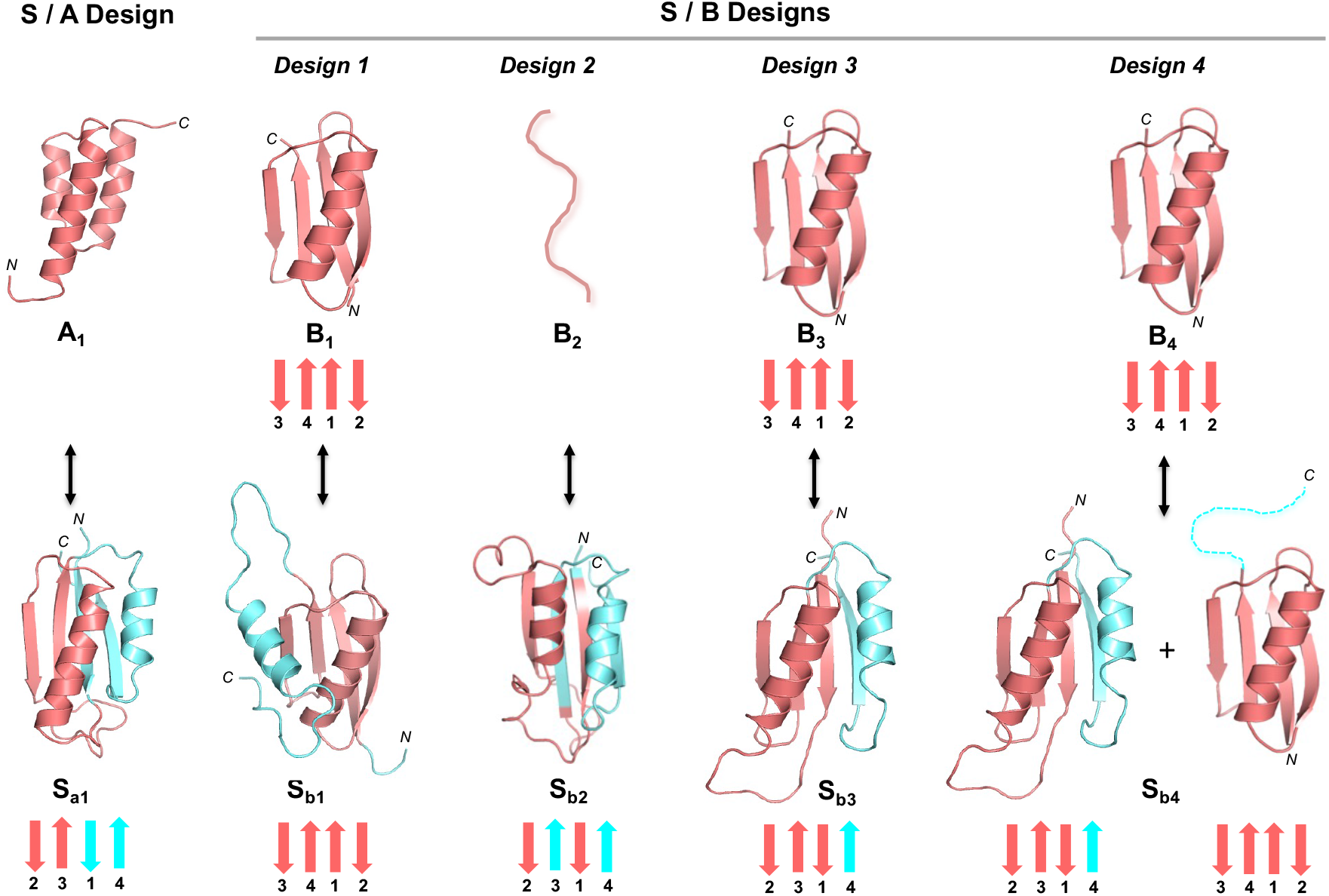
Summary of designed dimorphic proteins. Cartoon structures of five pairs of dimorphic proteins are depicted based on the NMR structures of A_1_, B_1_, B_4_, S_a1_, S_b1_, S_b2_, and S_b3_ that are determined here. The common sequence in each pair is in red. The extra amino acids in the longer sequences are in cyan. The arrows at the bottom of β-sheet containing structures show the topology of β−strands.

## Results

### Design of S_a1_ and A_1_

Designing a dimorphic sequence is an iterative process. After examining the 40 possible alignments of the 56 amino acid A-sequence in the 95 amino acid S-fold, we chose amino acids 11-66 as the preferred region of overlap (**Fig. S2**). This alignment generated nine positions of identity between the starting sequences. In terms of topological alignment, α1(S) mostly coincides with α1(A). β2(S) becomes α2(A), and the long β2-β3 turn and first half of β3(S) becomes α3(A). β1 and α2-β4 of the S-fold are outside of the overlap region. Mutations to resolve catastrophic interactions in this alignment were designed in clusters of 4-6 amino acids using Pymol (38) and relaxed structures were generated using Rosetta-Relax (35). Ultimately, we made 25 substitutions of an S-residue with an A-residue, substituted with a third choice in 5 cases, and retained the S-residue at 26 positions. We then examined the non-overlapping region of sequence and made 14 additional mutations to generate the S_a1_ sequence. The 56 amino acid version of the protein has 22 total changes: 17 substitutions of an A-with the S-residue and 5 changes to a third choice (**Fig. S2**). The final computational models for S_a1_ and A_1_ were generated by Rosetta using the Relax application. The Relax protocol searches the local conformational space around the native, experimentally-determined structure and is used only to evaluate whether the designed mutations have favorable native interactions within that limited conformational space. The designed models of S_a1_ and A_1_ show relatively small increases in energy compared to the relaxed native structures (Supplemental PDB files of the Rosetta models).

### Structure of A_1_

Overall, the 3α-helical bundle topology of A_1_ is very similar to the G_A_ parent structure from which it was derived (39). The sequence specific chemical shift assignments for A_1_ (**Fig. 2A**) were utilized to calculate a 3D structure with CS-Rosetta (**Fig. 2B, Table S1**). Our previous studies indicated close correspondence of CS-Rosetta and *de novo* structures for A- and B-folds (40). The N-terminal residues 1-4 and the C-terminal residues 53-56 are disordered in the structure, consistent with {^1^H}-^15^N steady state heteronuclear NOE data (**Fig. 2D**).

**Figure 2:**
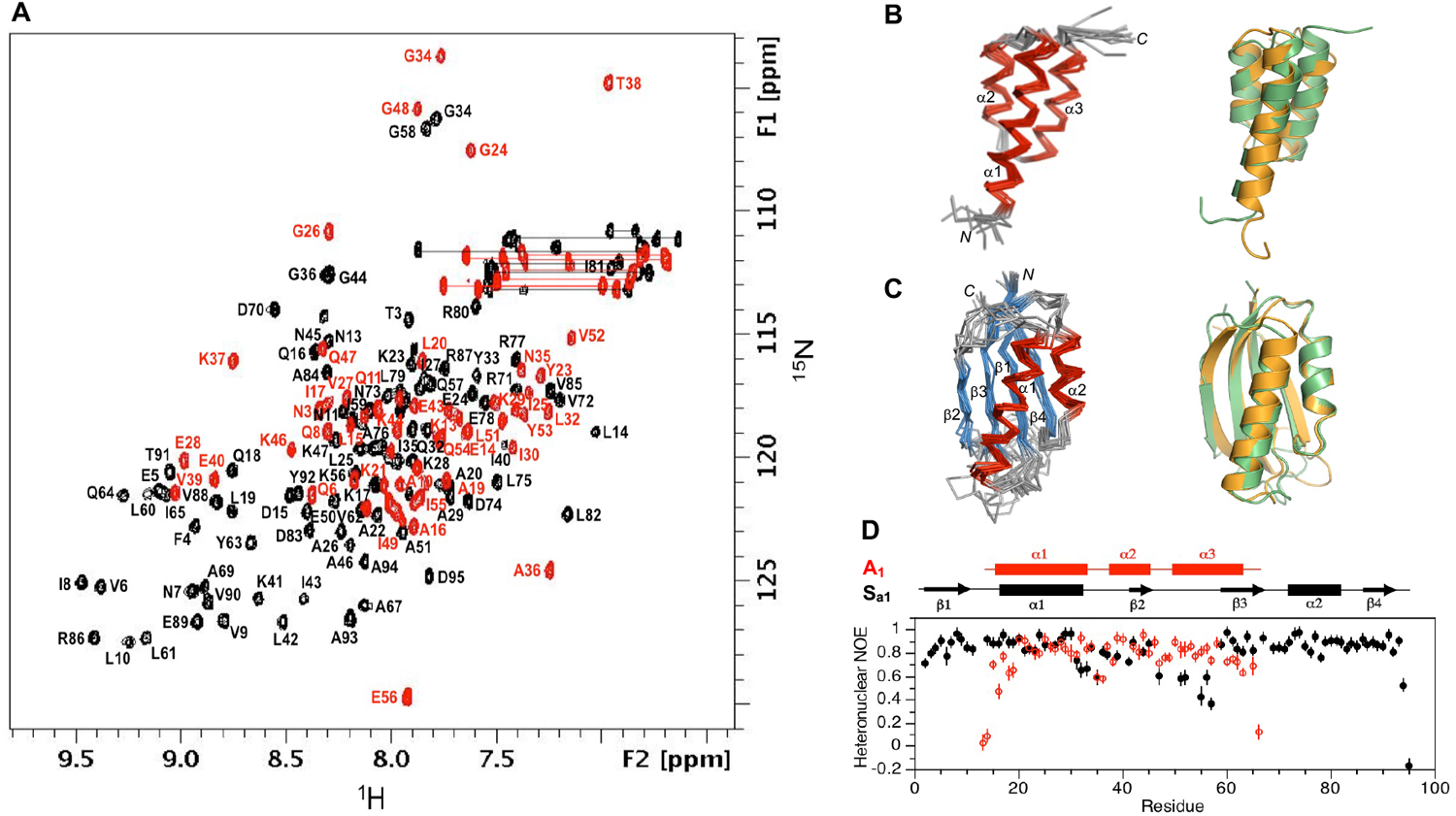
Structure and dynamics of A_1_ and S_a1_. (**A**) Overlaid two dimensional ^1^H-^15^N HSQC spectra of S_a1_ (black) and A_1_ (red) with backbone amide assignments. Spectra were recorded at 298°K and 278°K, respectively. (**B**) Ensemble of 10 lowest energy CS-Rosetta structures for A_1_ (left panel). Superposition of the A_1_ structure (green) with the parent G_A_ fold (orange) (right panel). (**C**) Ensemble of 10 lowest energy CS-Rosetta structures for S_a1_ (left panel). Superposition of S_a1_ (green) with the parent S6 fold (orange) (right panel). (**D**) Backbone dynamics in designed proteins. Plot of {^1^H}-^15^N steady state heteronuclear NOE values at 600 MHz versus residue for A_1_ (red) and for S_a1_ (black). Error bars indicate ±1SD.

### Structure of S_a1_

Likewise, S_a1_ has the same overall βαββαβ-topology as the parent S6 structure (**Fig. 2C, Table S2**). The backbone chemical shifts (**Fig. 2A**) were used in combination with main chain inter-proton NOEs (**Fig. S3**) to determine a three-dimensional structure using CS-Rosetta. The conformational ensemble shows well-defined elements of secondary structure at residues 2-10 (β1), 16-32 (α1), 40-44 (β2), 59-67 (β3), 73-81 (α2) and 86-92 (β4). The principal difference from the native structure is that the β2-strand is seven amino acids shorter in S_a1_ than in S6. Heteronuclear NOE data show overall consistency with the structure, indicating that the loop between the β2- and β3-strands from residues 45-58 is more flexible than other internal regions of the polypeptide chain (**Fig. 2D**).

Although the 56 amino acid sequence of A_1_ is 100% identical to residues 11-66 of S_a1_, a significant fraction of the residues undergo large amplitude changes in their backbone ϕ/ψ torsion angles between these two structures (**Fig. 3A**). Amino acids 1-4 form the disordered N-terminal tail in A_1_ and are part of the loop between the β1-strand and α1-helix in S_a1_ (residues 11-14). Amino acids 5-23 form the α1-helix in A_1_ and the equivalent sequence in S_a1_ (residues 15-33) forms a similar length α1-helix. Amino acids 24-26 form the loop between the α1- and α2-helices in A_1_ and form the first part of the loop between the α1-helix and β2-strand in S_a1_ (residues 34-36). Amino acids 27-35 in the α2-helix of A_1_ correspond with the extended part of the α1-β2 loop and the β2 strand in S_a1_ (residues 37-45). Amino acids 36-38 form the loop between the α2- and α3-helices in A_1_ and are part of the loop between the β2- and β3-strands in S_a1_ (residues 46-48). Amino acids 39-56 in the α3-helix and C-terminal tail of A_1_ form a portion of the β2-β3 loop and the β3-strand in S_a1_ (residues 49-66).

**Figure 3:**
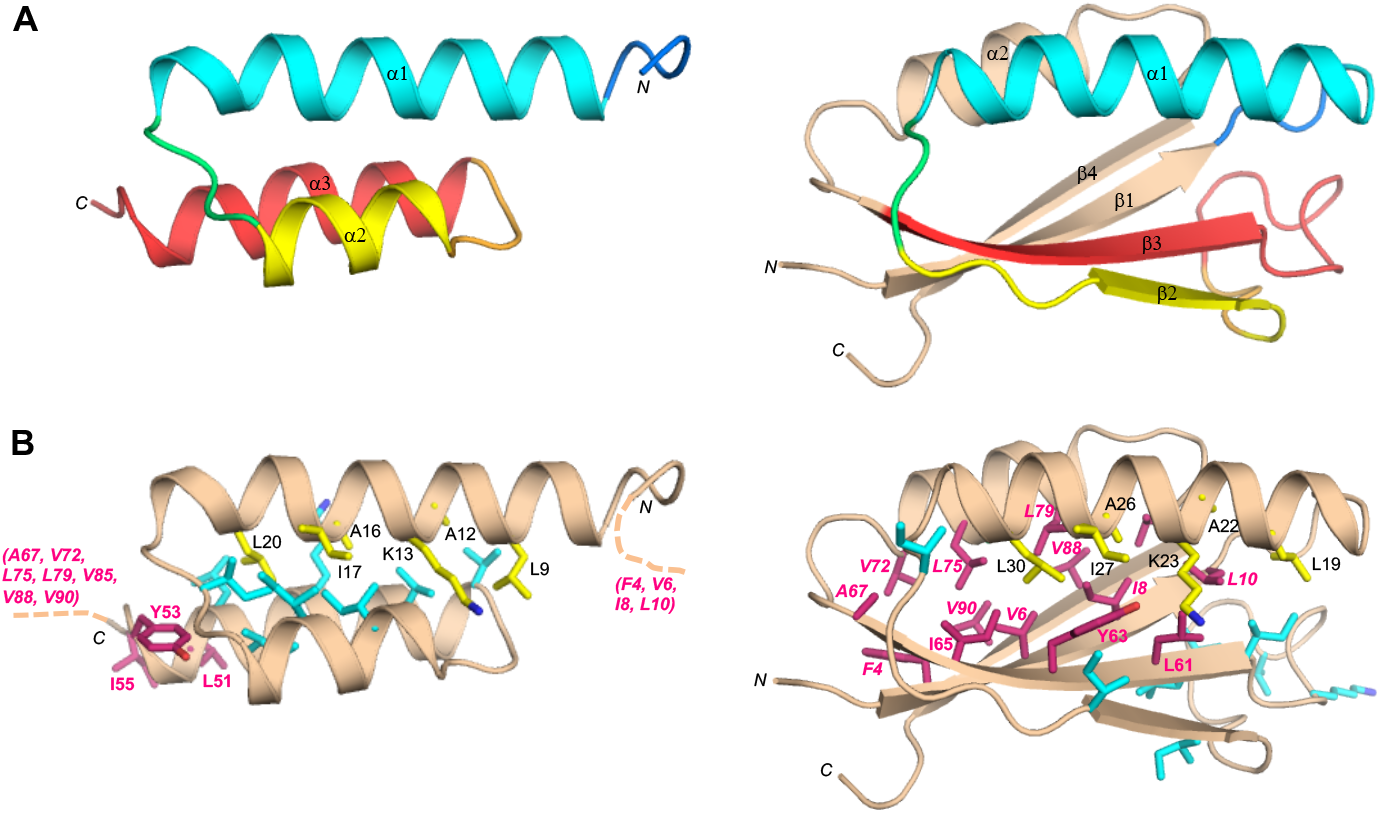
Structural differences between the sequence identical regions of A_1_ and S_a1_. (**A**) Main chain comparisons. (Left panel) CS-Rosetta structure of A_1_ with color coding for secondary structured elements. (Right panel) Corresponding color-coded regions mapped onto the CS-Rosetta structure of S_a1_, illustrating changes in backbone conformation. Regions outside the 56 amino acid sequence of A_1_ are shown in wheat. (**B**) Side chain comparisons. (Left panel) Residues contributing to the core of A_1_ from the α1-helix (yellow), and from other regions (cyan). The non-α1 core residues from S_a1_ (pink) do not overlap with the A_1_ core (see text for further details). (Right panel) Residues contributing to the core of S_a1_ from the α1-helix (yellow), and most of the other participating core residues (pink). The non-α1 core residues from A_1_ are also shown (cyan), highlighting the low degree of overlap.

The CS-Rosetta structures calculated here employ main chain chemical shift and NOE restraints but do not have experimental restraints for side chains. Nevertheless, the overall positions of the core side chains are very likely to be correct to a close approximation due to the packing requirements dictated by the respective folds. It is therefore instructive to compare the location of corresponding modeled side chains for core residues in the 3α versus α/β-plait folds of these NMR-derived structures. Residues contributing to the core in A_1_ consist of L9, A12, K13, A16, I17, L20, and Y23 from the α1-helix; I25 from the loop between the α1- and α2-helices; I30 and I33 from the α2-helix; and V39, V42, K46, I49, and L50 from the α3-helix (**Fig. 3B**). The core of S_a1_ is somewhat larger with 21 residues versus 15 residues for A_1_. Core amino acids in the α1-helix of A_1_ correspond with residues that also contribute to the core of S_a1_. Only two of these amino acids, A16/A26 and L20/L30, are completely buried in both folds. In contrast with the 3α fold of A_1_, the α1-helix in the α/β-plait fold of S_a1_ contacts an almost entirely different set of residues. For example, amino acids L51, Y53, and I55 in the C-terminal tail of A_1_ do not have extensive contacts with α1 but the corresponding residues in S_a1_ (L61, Y63, and I65) form close core interactions with α1 as part of the β3-strand. Most of the other core residues contacting the α1-helix of S_a1_ are outside the 56 amino acid region coding for the A_1_ fold. These include F4, V6, I8, and L10 from the β1-strand; A67 from the β3-strand; V72, L75, and L79 from the α2-helix; and V85 from the loop between the α2-helix and the β4-strand. Two additional residues, V88 and V90 (β4) also contribute significantly to the core but do not contact α1. Thus, beyond the original topological alignment of the α1-helices, the cores of the 3α and α/β-plait folds are largely non-overlapping. In total, approximately half (11/21) of the residues participating in the S_a1_ core are not present in the A_1_ sequence. This includes residues 1-10 at the N-terminus, which contribute 4 amino acids to the S_a1_ core (F4, V6, I8, L10), and residues 67-95 which provide 7 core amino acids (A67, V72, L75, L79, V85, V88, and V90).

### CD analysis of unfolding for A_1_/S_a1_

Far-UV CD spectra were measured for S_a1_ and A_1_ and thermal unfolding profiles were determined by measuring ellipticity at 222nm vs. temperature (**Fig. S4**). The fraction native was determined by subtracting an unfolded baseline from the experimental CD signal and then dividing by the total CD difference between 100% folded and 0% folded at that temperature. Reversibility of unfolding was confirmed by comparing the CD spectra at 293^∘^K before melting and after heating to 373^∘^K and cooling to 293^∘^K. The temperature unfolding profiles were converted to an apparent ΔGfolding and fit to a theoretical curve calculated using the Gibbs-Helmholtz equation: ΔGfolding = ΔHo-TΔSo +ΔCp(T-To-TlnT/To), where To = 298^∘^K (**Fig. 4A, B**) (41). S_a1_ has a T_M_ of ∼373^∘^K and an estimated ΔGfolding of −7.5 kcal/mol at 298^∘^K (**Fig. 4B, C**). The ΔG_folding_ of the parent S6 is −8.5 kcal/mol (32). The Rosetta energy of the S_a1_ design is similar to the native sequence (**Fig. 4D**). A_1_ has a ΔG_folding_ = −4.0 kcal/mol at 298^∘^K. The ΔG_folding_ of the parent is −5.6 kcal/mol (42, 43). The Rosetta energy of the A_1_ design is slightly more favorable than the native sequence (**Fig. 4D**).

**Figure 4:**
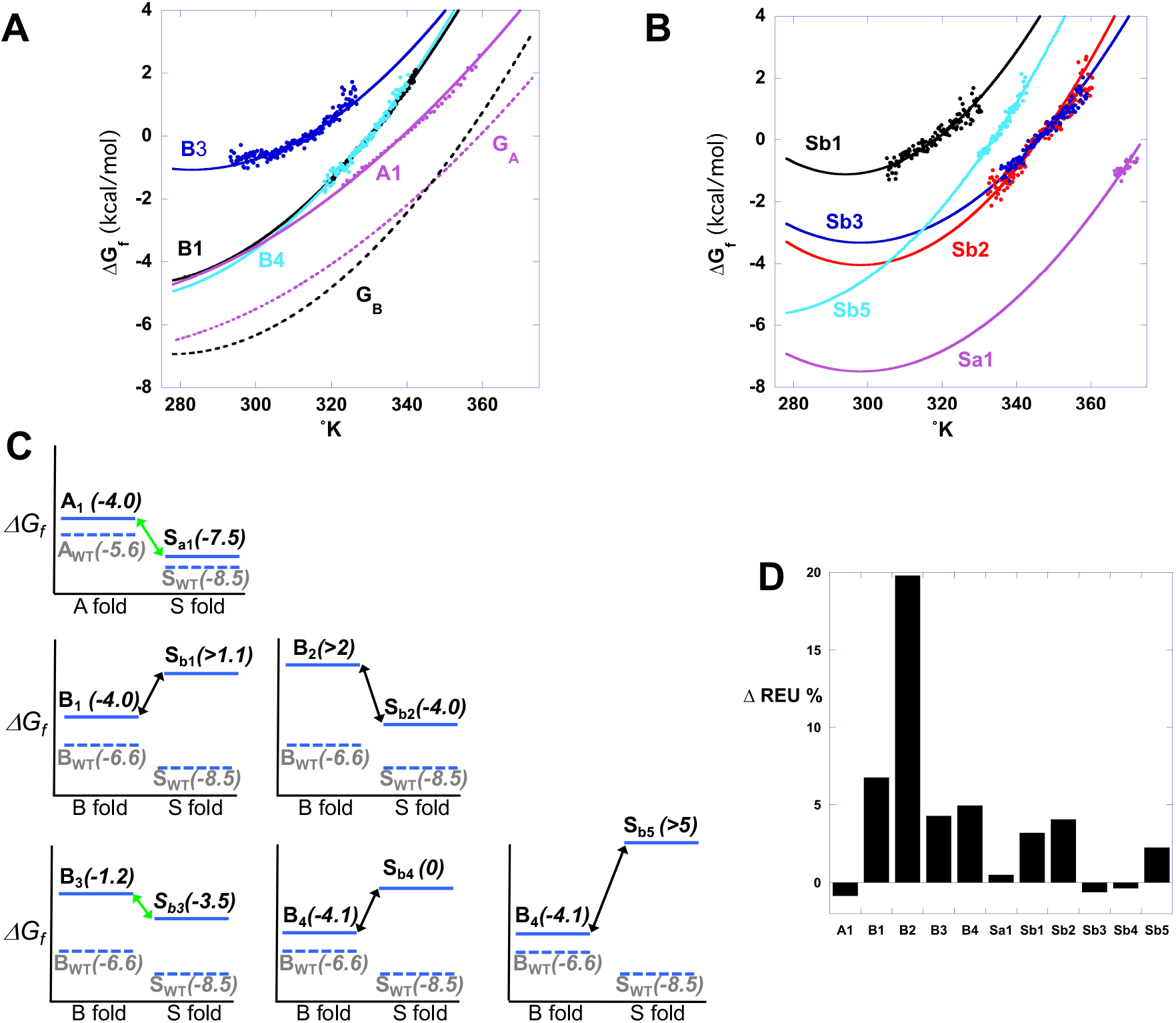
Energetics of fold switches. **(A)** ΔG vs T profiles for 56 residue proteins. **(B)** ΔG vs T profiles for longer proteins. ΔG_folding_ is plotted vs temperature in order to assess stability at a reference state of 298^∘^K. The curvature of the profiles reflects the ΔCp of folding for each protein (41). ΔCp = −0.69 kcal/°mol for G_B_ and −0.26 kcal/°mol for G_A_. (42, 44) and −1.1 kcal/°mol for S6. **(C)** Each panel shows the ΔG_folding_ for the 56 residue G-fold and longer S-fold in a dimorphic pair. For example, the 56 residue A_1_ protein has a ΔG_folding_ of −4.0 kcal/mol for the A-fold. The same 56 residues in the longer S_a1_ protein has a ΔG_folding_ of −7.5 kcal/mol for the S-fold. The green connecting arrows indicate a complete fold switch for these sequence pairs. **(D)** The percent change in Rosetta Energy Units (REU) between the parent protein and the designed switch protein is plotted. The computational design of A_1_ was compared to the relaxed structure of a highly stable A-fold (43). The designs of B_1_, B_2_, B_3_, and B_4_ were compared with a highly stable B-fold (44). The designs of S_a1_, S_b1_, S_b2_, S_b3_, S_b4_, and S_b5_ were compared to a highly stable S-fold (32). All designed proteins have relatively small changes in REU except for B_2_. The 20% increase in REU for B_2_ is consistent with its low stability.

### Design 1 of S_b1_ and B_1_

After examining the 40 possible alignments of the 56 residue B-sequence in the 95 residue S-fold, we chose amino acids 4-59 as the preferred region of overlap (**Fig. S5**). This alignment generated five positions of identity between the starting sequences. In terms of topological alignment, β1(S) mostly coincides with β1(B). The first half of α1(S) becomes β2(B) and the second half of α1(S) becomes the first half of α1(B). The β2 strand ofS becomes the second half of α1(B), a turn, and the first part of β3(B). The long β2-β3(S) turn and the first part of β3(S) become the second part of β3 and β4 of B. The second half of β3 and α2-β4 of the S-fold are outside of the overlap region. At the 51 positions of non-identity from 4 to 59, we made 40 substitutions of an S-residue with an A-residue, substituted with a third choice in 4 cases, and retained the S-amino acid at 7 positions. We then made 14 additional mutations in the non-overlapping region to generate the S_b1_ sequence. The 56 amino acid version of the protein has 11 total changes: 7 substitutions of an A-residue with the S-residue and 4 changes to a third choice (**Fig. S5**). The energies of the computational models for S_b1_ and B_1_ show relatively small increases in energy compared to the relaxed native structures (**Fig. 4D**).

### Structure of B_1_

The ββαββ topology of B_1_ is very similar to that of the parent B-fold, with a backbone RMSD of ∼0.6Å. The NMR structure consists of four β-strands defined by residues 2-9 (β1), residues 13-20 (β2), residues 42-46 (β3), and residues 50-55 (β4) and one α-helix from residues 23-37 (**Fig. 5A, Fig. S6B, Table S1**).

**Figure 5:**
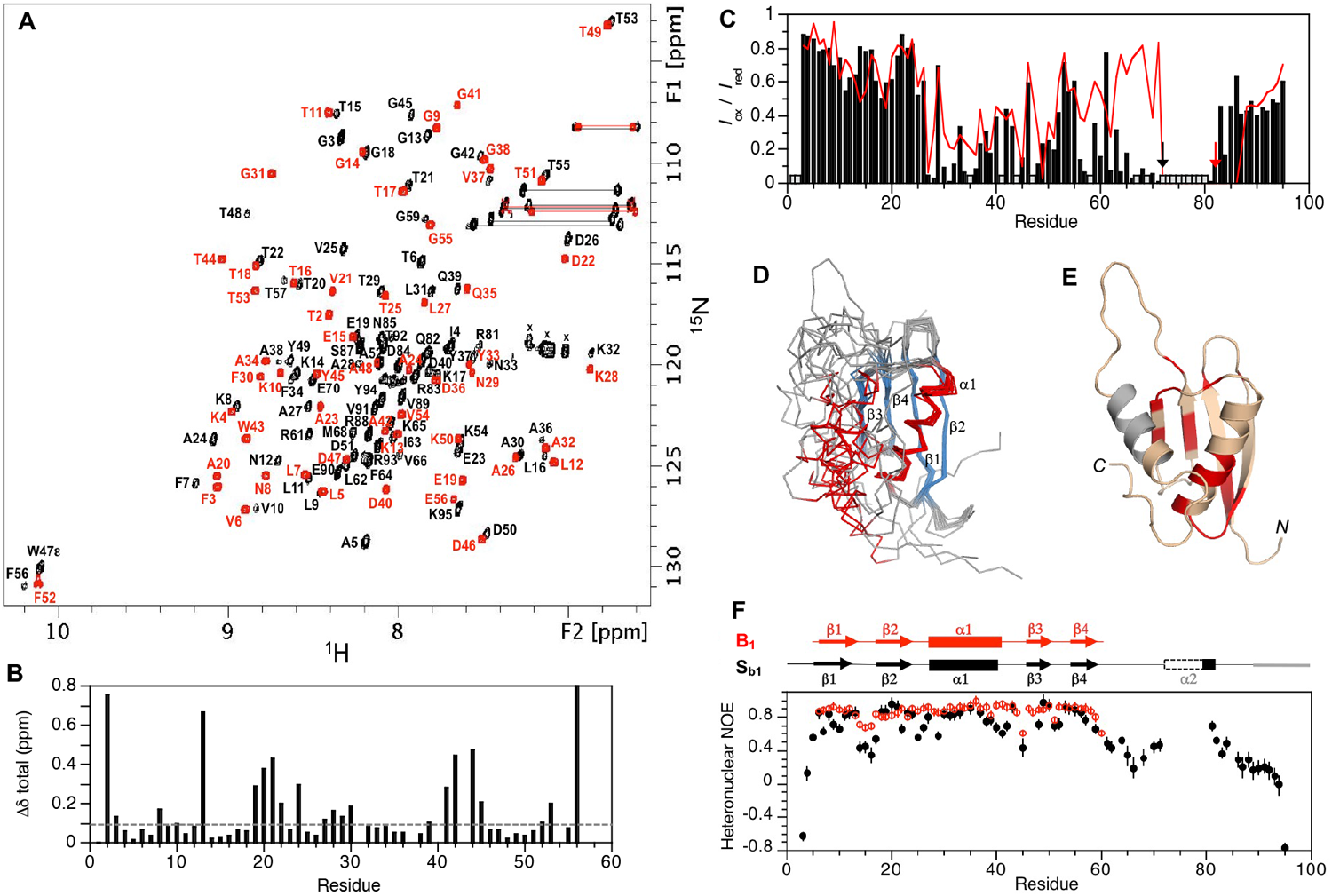
Comparison of B_1_ and S_b1_. (**A**) Overlaid two dimensional ^1^H-^15^N HSQC spectra of S_b1_ (black) and B_1_ (red) with backbone amide assignments. Spectra were recorded at 283°K. (**B**) Plot of chemical shift perturbations between S_b1_ and B_1_ for backbone amides in the 56 amino acid identical region. Residue numbering is for B_1_. (**C**) Plot of I_ox_/I_red_ versus residue for S_b1_-R72C-MTSL (black) and S_b1_-R83C-MTSL (red). Gray columns indicate unassigned residues or prolines. The positions of the spin labels are indicated with arrows. (**D**) Ensemble of 10 lowest energy CS-Rosetta structures for S_b1_ using PRE restraints. (**E**) Cartoon representation of model 1 from the ensemble. Values of βδ_total_>0.1 ppm from (B) are mapped onto the structure (red). Unassigned residues in the putative α2-helix are in gray. (**F**) Plot of {^1^H}-^15^N steady state heteronuclear NOE values at 600 MHz versus residue for B_1_ (red) and for S_b1_ (black). Error bars indicate ±1SD.

### Structure of S_b1_

The topology of S_b1_ is not the same as the parent S6 structure. Instead, the 2D ^1^H-^15^N HSQC spectrum of S_b1_ has a pattern similar to that of B_1_ (**Fig. 5A**). NMR assignment of the main chain resonances showed the presence of four β-strands and two α-helices, but the order of the secondary structure elements was ββαββα rather than the βαββαβ arrangement expected for an S-type fold. Initial NMR structures of S_b1_ indicated a B-fold, which was supported by backbone NOE connectivities (**Fig. S3**), with a mostly disordered C-terminal tail. CS-Rosetta modeled residues 73-83 near the C-terminus as an α2-helix. Of these, amide signals due to residues 73-80 were not detectable in NMR spectra while residues 81-83 were helical based on assigned chemical shifts. Comparison of S_b1_ amide chemical shifts with those of B_1_ indicated that most of the perturbations due to the C-terminal 35 amino acid tail were localized in α1, β3, and neighboring regions (**Fig. 5B**,**E**). This suggested that the putative α2-helix interacts with the B-fold in these contiguous regions. Mutations R72C and R83C were made at the N- and C-terminal ends of the α2 region in separate samples of S_b1_ and these proteins were derivatized with the stable nitroxide spin label MTSL. Paramagnetic relaxation enhancement (PRE) measurements (**Fig. 5C**) showed significant decreases in amide peak intensity over the α1 and β3 regions for the B-core of S_b1_, consistent with the chemical shift perturbation data. Furthermore, the PRE intensity profiles were similar regardless of which end of the α2 region the spin label resided. This suggests that docking of the α2 region against the B-folded core of S_b1_ is in exchange between multiple states, providing a plausible explanation for why most of the α2 amide resonances are not detectable. Structures for S_b1_ were re-calculated using additional weak (<20Å) PRE restraints, showing an ensemble with a well-defined B-core that has a putative α2-helix packed against it loosely (**Fig. 5D, E**). Steady-state {^1^H}-^15^N heteronuclear NOE data for S_b1_ were consistent with the structure (**Fig. 5F**). In particular, the C-terminal tail becomes more ordered around the α2 region, although these heteronuclear NOE values (0.4-0.7) are still below those of well-ordered regions (>0.8). Thus, the structure of S_b1_ may be viewed as a transitory state between the S- and B-folds. With two α-helices packed against a 4-stranded β-sheet, S_b1_ has the same overall two-layer α/β-sandwich architecture as the S-fold but differs in the topological arrangement of secondary structures.

### CD analysis of unfolding for B_1_/S_b1_

B_1_ has a ΔG_folding_ = −4.0 kcal/mol at 298^∘^K compared to −6.6 kcal/mol for the parent (44). The Rosetta energy of the B_1_ design is a little less favorable than for the native sequence and generally consistent with its ΔG_folding_ (**Fig. 4C, D**). In contrast, S_b1_ has a minimum ΔG_folding_ = −1.1 kcal/mol at ∼298^∘^K (**Fig. 4B**). As described above, the predominant folded form at 298^∘^K is the B-conformation. The energy of its Rosetta design in the S-conformation is similar to that of S_a1_, however. Thus, while B_1_ and S_b1_ have identical sequences in their respective 56 amino acid B-folds, the 35 residue C-terminal tail in S_b1_ destabilizes the B-fold, presumably by populating competing alternative states. Elucidating the reason for the large inconsistency between the S_b1_ design energy and the observed fold is critical for switch design and was further investigated below.

### Design 2 of S_b2_ and B_2_

We introduced 13 mutations into the S_b1_ sequence to generate a second dimorphic version in this alignment (**Fig. S5**). The Rosetta energy of the S_b2_ design model is almost identical to the S_b1_ design model. The 56 amino acid version of S_b2_ (denoted B_2_) has a significantly higher Rosetta energy than B_1_ (**Fig. 4D**), however.

### Structure of S_b2_

The 3D structure of S_b2_ contains four β-strands and two α-helices and has the general features of the parent S-fold (**Fig. S6, Table S2**). The ordered regions in the structure are residues 1-9 (β1), 23-32 (α1), 43-48 (β2), 59-65 (β3), 71-80 (α2), and 86-91 (β4). While the parent S (PDB 1RIS) and S_a1_ structures are very similar, the S_b2_ structure differs from both in a number of ways despite having the same overall topology. The α1-helix in S_b2_ is shorter, comprising 10 amino acids compared with 17 amino acids in S_a1_. Also, the β2-strand forms 4 amino acids later in the S_b2_ polypeptide chain than in S_a1_. The first residue in the β2-strand of S_b2_, G43, interacts with E65 in the β3-strand. This represents a two-residue shift in the register of hydrogen bonding between β2 and β3 in S_b2_ compared with S_a1_ (**Fig. S3B**). As a result of these differences, the loops connecting β1 to α1 and α1 to β2 are longer in S_b2_ (13 and 10 residues, respectively) than in S_a1_ (5 and 7 residues, respectively). The remainder of the S_b2_ structure encompassing β1, β3, α2, and β4 is very similar to S_a1_. Heteronuclear NOE dynamics data for S_b2_ were consistent with the NMR structure (**Fig. S6D**). In particular, the relatively long β1-α1, α1-β2, and β2-β3 loops were found to be the most flexible on the ns-ps timescale. We were not able to characterize the B_2_ structure because it was largely unfolded, consistent with its increased Rosetta energy. Instead, the structure of B_1_ was used for comparisons with S_b2_ because the corresponding B-regions also have very high sequence identity (80%). Detailed comparisons between B_1_ and S_b2_ are presented in Supplemental Material (**Fig. S7**, and Supplementary Information).

### CD analysis of unfolding for B_2_/S_b2_

S_b2_ has a minimum ΔG_folding_ = −4.0 kcal/mol at ∼298^∘^K (**Fig. 4B**). As described above, the predominant folded form at 298^∘^K is the S-conformation. Its Rosetta design energy is actually slightly less favorable than the design energy for S_b1_, however. From CD, B_2_ appears to be ≥95% unfolded conformation (ΔG_folding_ ≥2kcal/mol) throughout the temperature range from 278-373^∘^K, consistent with its unfavorable Rosetta energy (**Fig. 4D**). Thus the fold switch between S_b1_ (B-fold) and S_b2_ (S-fold) appears to result from decreased stability of the embedded B-fold rather than improved interactions in the folded S-conformation.

### Design 3 of S_b3_ and B_3_

Analysis of the NMR structures of S_b1_ and S_b2_ provided clues about how to improve the design of dimorphic B-fold/S-fold proteins. In the computational design of S_b1_, the DDATK turn should become part of the long connection between β2 and β3 of the S-fold. The sequence actually remains in the B-conformation, however. This occurs in spite of acceptable native interactions in the S-conformation, as assessed by Rosetta. We did not anticipate that turn propensities would be harder to override than secondary structure propensities but clearly turn sequences (even without proline or glycine) can contain critical topological information (45-48). The S_b2_ sequence had two substitutions in the DDATK sequence which decrease its strong propensity for the short turn. Based on this insight, we redesigned the S-fold to increase its compatibility with the B-fold. The S-fold is classified as a superfold with many natural variations in the length and position of turns (49). In particular, some natural S-folds (protease inhibitors) have a short turn between β2 and β3 that matches the B-fold turn between β3 and β4 (50-52). These protease inhibitors have a longer loop between β1 and α1. We made an S-fold of this type by inserting three residues (GTD) between β1 and α1 and deleting 12 residues (RQLSEPIAKDPQ) from the long loop between β2 and β3 (**Fig. S8, S9**). This creates a topological match between α1β3β4 in B and α1β2β3 in S.

In this design, amino acids 1-56 are the region of overlap (**Fig. S8**). In terms of topological alignment, the β1−strand is the same in both folds but changes orientation, the long turn between β1 and α1 in S_b3_ becomes β2 of the B-fold, and the α1−β2−β3 of S_b3_ maintains the same topology in the B_3_ design. The α2-helix and the β4−strand of S_b3_ are outside the overlap region. At the 47 positions of non-identity, we made 33 substitutions of an S-residue with a B-residue, substituted with a third choice in 12 cases, and retained the S-amino acid at one position. We then made 18 additional mutations in the non-overlapping region to generate the S_b3_ sequence. The 56 amino acid version of the protein has 12 total changes: 1 substitution of a B-residue with the S-residue and 11 changes to a third choice. The energy of the computational model for S_b3_ is slightly more favorable than the relaxed native structure. The designed model for B_3_ has a less favorable energy than the native B-sequence but is more favorable than the relaxed B_1_ design (**Fig. 4D**).

### Structural analysis of B_3_

The 2D ^1^H-^15^N HSQC spectrum of B_3_ at 278°K and low concentrations (<20 μM) was consistent with a predominant, monomeric B-fold (**Fig. S10**) but showed significant exchange broadening at 298°K, indicative of low stability (see below). Presumably the low stability is due to less favorable packing of Y5 in the core of the B-fold compared with a smaller aliphatic leucine. However, additional, putatively oligomeric, species were also present for which relative peak intensities increased with increasing protein concentration. Due to its relatively low stability and sample heterogeneity, B_3_ was not analyzed further structurally.

### Structural analysis of S_b3_

In contrast, when the 56-residue B_3_ sequence was embedded in the longer 87-residue polypeptide chain to give S_b3_, it provided a homogeneous sample for which the HSQC spectrum was readily assigned (**Fig. 6A**). NMR-based structure determination indicated that S_b3_ has a βαββαβ secondary structure and an S-fold topology (**Fig. 6C**). Ordered regions correspond with residues 4-10 (β1), 24-37 (α1), 42-46 (β2), 51-56 (β3), 62-70 (α2), and 79-85 (β4). Comparison of S_b3_ with the parent S-fold indicates that the β1/α2β4 portion of the fold is similar in both. In contrast, the β1-α1 loop is longer in S_b3_ (13 residues) than in the parent S-fold (5 residues), while α1, β2, the β2-β3 loop, and β3 are all shorter than in the parent (**Fig. 6C**). Consistent with the S_b3_ structure, the 13 amino acid β1-α1 loop is highly flexible (**Fig. 6D**).

**Figure 6:**
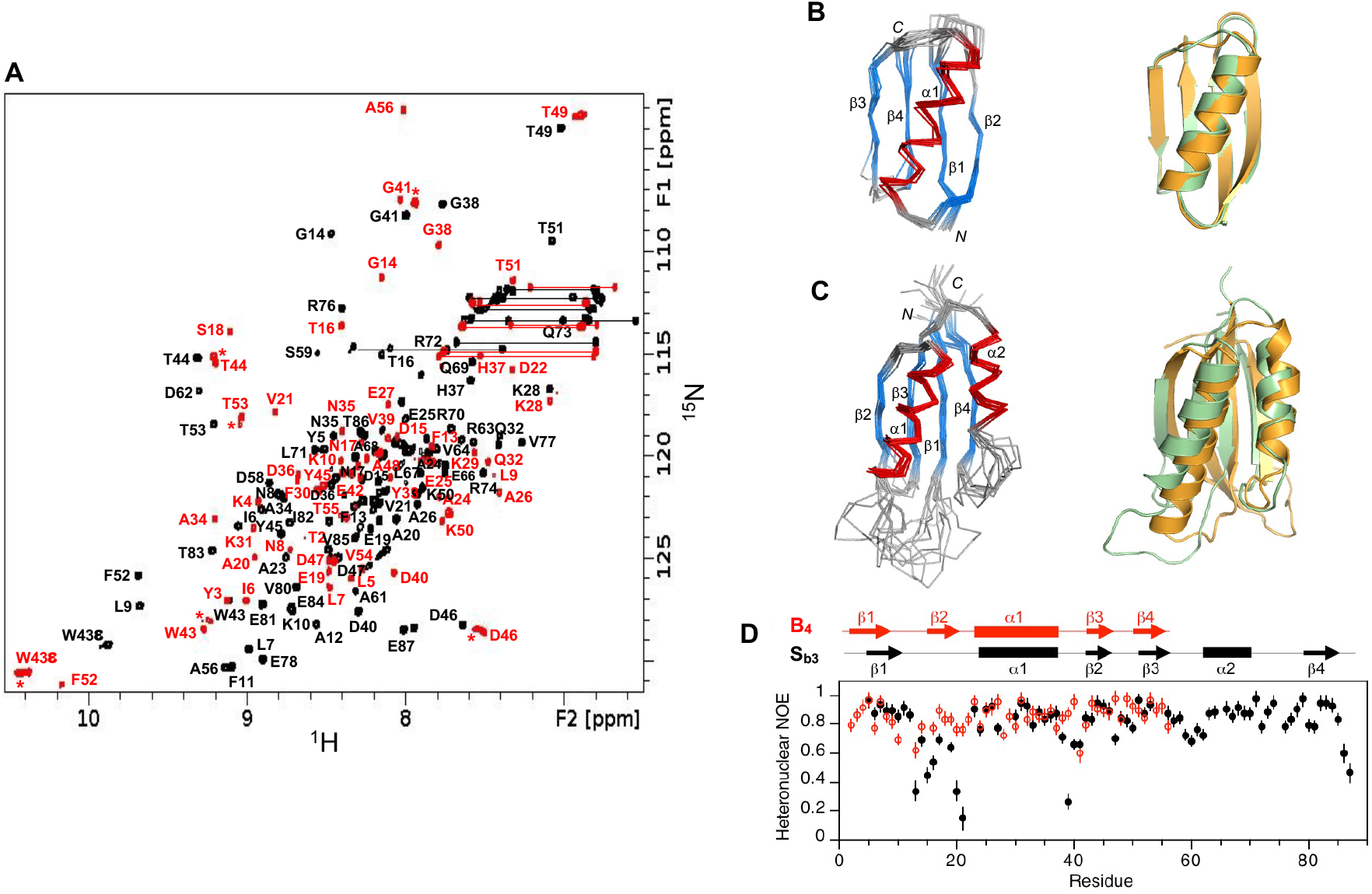
Structure and dynamics of S_b3_ and B_4_. (**A**) Overlaid two dimensional ^1^H-^15^N HSQC spectra of S_b3_ (black) and B_4_ (red) with backbone amide assignments. Spectra were recorded at 298°K. The A56 peak is an aliased signal. Peaks labeled with an asterisk decrease in relative intensity as the B_4_ concentration is lowered, indicating the presence of a weakly associated putative dimer in addition to monomer. (**B**) Ensemble of 10 lowest energy CS-Rosetta structures for B_4_ (left panel). Superposition of the designed B_4_ structure (green) with the parent G_B_ fold (orange) (right panel). (**C**) Ensemble of 10 lowest energy CS-Rosetta structures for S_b3_ (left panel). Superposition of S_b3_ (green) with the parent S6 fold (orange) (right panel). (**D**) Plot of {^1^H}-^15^N steady state heteronuclear NOE values at 600 MHz versus residue for B_4_ (red) and S_b3_ (black). Error bars indicate ±1SD.

### CD analysis of unfolding for B_3_/S_b3_

S_b3_ has a minimum ΔG_folding_ = −3.5 kcal/mol at ∼298^∘^K (**Fig. 4B,C**). As described above, the predominant form is an S-fold. The Rosetta energy of its design is slightly more favorable than the energy of the S_a1_ design even though the stability of its S-fold is less by ∼4 kcal/mol (**Fig. 4C,D**). B_3_ has a ΔG_folding_ of −1.2 kcal/mol at 298^∘^K (**Fig. 4A,C**). The ΔG_folding_ of B_3_ is less than would be expected from its Rosetta energy. From the NMR analysis, it appears that the B-fold is in equilibrium with putatively dimeric states. This creates a situation in which the B-fold is both temperature dependent and concentration dependent. The predominant form at 278^∘^K and ≤18µM is the B-fold, however. The low stability and concentration-dependent behavior of B_3_ may indicate that some propensity for the S-conformation could persist in the 56-residue protein.

### Design of S_b4_ and B_4_

We used the NMR structure of S_b3_ to design a point mutation (Y5L) that would stabilize the embedded B-fold and simultaneously destabilize the S-fold. This was expected to shift the population to the B-fold. The Y5L mutation was also introduced into B_3_ to determine its effect on the stability of the B-fold in the 56 amino acid protein. These new dimorphic proteins are denoted S_b4_ and B_4_.

### Structural analysis of B_4_

Assignment and structure determination of B_4_ showed its topology to be identical to the parent B-topology (**Fig. 6A, B**). At concentrations above 100 μM, B_4_ displayed a tendency for weak self-association similar to that seen for B_3_.

### Structural analysis of S_b4_

Incorporation of the single amino acid change Y5L into S_b3_ to give S_b4_ resulted in approximately twice the number of amide cross-peaks in the HSQC spectrum relative to the S_b3_ sample. Comparison of the spectrum of S_b4_ with spectra of S- and B-folds for the closely related sequences of B_4_ and S_b3_ indicated that S_b4_ populates both S- and B-states simultaneously in an approximately 1:1 ratio at 298°K (**Fig. S11**). Due to this heterogeneity, the structure of S_b4_ was not analyzed further here.

### Comparison of S_b3_ and B_4_

The aligned amino acid sequences of S_b3_ and B_4_ show that their B-regions have 98% sequence identity (**Fig. S8**), the only difference being an L5Y mutation in S_b3_. The global folds of S_b3_ and B_4_ have large-scale differences, however (**Fig. 7A**). The β1-strands, while similar in length, are in opposite directions in S_b3_ and B_4_. The β1-strand forms a parallel stranded interaction with β4 in B_4_, but an antiparallel interaction with the corresponding β3-strand in S_b3_. Whereas residues 9-20 form the 6-residue β1-β2 turn and the 6-residue β2-strand of B_4_, these amino acids constitute the end of β1 and 10 residues of the large disordered β1-α1 loop in S_b3_. The remainder of the B-region is topologically similar, with the α1/β3/β4 structure in B_4_ matching the α1/β2/β3 structure in S_b3_. Overall, however, the order of H-bonding in the 4-stranded β-sheets is quite different, with β2β3β1β4 in S_b3_ and β3β4β1β2 in B_4_.

**Figure 7:**
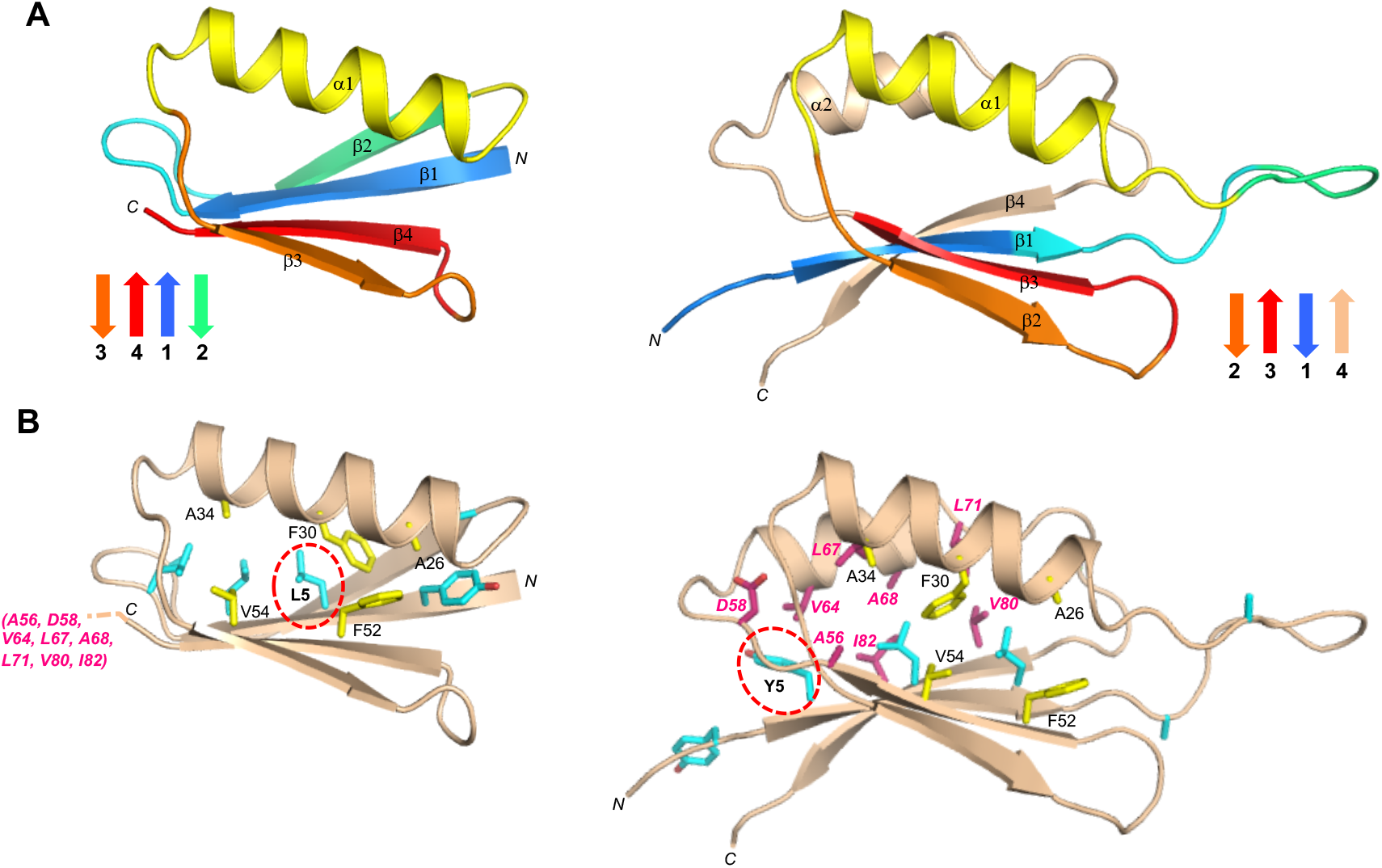
Structural differences in the high sequence identity regions of B_4_ and S_b3_. (**A**) Main chain comparisons. (Left panel) CS-Rosetta structure of B_4_ with secondary structure elements color coded. (Right panel) Corresponding color-coded regions mapped onto the CS-Rosetta structure of S_b3_, showing changes in backbone conformation. Regions outside the 56 amino acid sequence of B_4_ are shown in wheat. (**B**) Side chain comparisons. (Left panel) Residues contributing to the core of B_4_ from α1/β3/β4 (yellow), and from other regions (cyan). The non-α1/β2/β3 core residues from S_b3_ (pink) do not overlap with the B_4_ core (see text for further details). (Right panel) Residues contributing to the core of S_b3_ from α1/β2/β3 (yellow), and the other participating core residues (pink). The non-α1/β2/β3 core residues from B_4_ are also shown (cyan). The single L5Y amino acid difference between B_4_ and S_b3_ is highlighted.

The main core residues of B_4_ consist of Y3, L5, L7, and L9 from β1, A26, F30, and A34 from α1, and F52 and V54 from β4 (**Fig. 7B**). In S_b3_, the topologically equivalent regions of the core are A26, F30, and A34 from α1, and F52 and V54 from β3. Residues Y5, L7, and L9 from the β1 strand of S_b3_ also form part of the core, but with different packing from B_4_ due to the reverse orientation of β1. Residues A12 and A20, which contribute to the periphery of the core in B_4_, are solvent accessible in the β1-α1 loop of S_b3_. Most of the remaining core residues of S_b3_ come from outside of the B-region and include amino acids from β3 (A56), α2 (V64, L67, A68, L71), and β4 (V80 and I82). Overall, the degree of overlap between the cores of B_4_ and S_b3_ is higher than for A_1_/S_a1_ and B_1_/S_b2_ (compare **Figs. 3, 6, and S7**), indicating that while mutual exclusivity of cores may be advantageous for fold switching it is not an absolute requirement.

### CD analysis of unfolding for B_4_/S_b4_

Thermal denaturation by CD shows that B_4_ has a ΔG_folding_ = −4.1 kcal/mol at 298^∘^K (**Fig. 4A, C**). The stability of S_b4_ can be derived from the NMR analysis because S- and B-folds are observed in equal mixture. Thus, the ΔG_folding_ for both the S- and B-folds of S_b4_ is ∼ 0 kcal/mol. Appending the 31 residue C-terminal tail of S_b4_ therefore destabilizes the B-fold by 4.1 kcal/mol and observably populates the S-fold. Notably, the Rosetta energy of the design model of S_b4_ is virtually identical to that of S_b3_, even though S_b3_ actually has a much more stable S-fold (**Fig. 4C**). This reflects the influence of the antagonistic B-fold on the S-fold population in S_b4_. The antagonism of the B-fold is not reflected in the Rosetta energy of S_b4_, however, because the Relax protocol examines only a limited conformational space around the design model.

### Design of S_b5_

We designed an L67R mutation in S_b4_ to destabilize the S-fold without changing the sequence of the embedded B-fold. The mutant is denoted as S_b5_. This was expected to shift the population to the B-fold. The energies of the computational models for S_b4_ and S_b5_ are shown in **Fig. 4D**. Note that the amino acid sequence for the 56 residue B-regions of S_b4_ and S_b5_ are the same as for B_4_.

### Structural analysis of S_b5_

The 2D ^1^H-^15^N HSQC spectrum of S_b5_ indicates that the L67R mutation does indeed destabilize the S-fold, with the loss of S-type amide cross-peaks and the concurrent appearance of a new set of signals. Superposition of the spectrum with that of B_4_ shows that the new signals in S_b5_ largely correspond with the spectrum of B_4_ (**Fig. S12**). Thus, the L67R mutation shifts the equilibrium from the S-fold to the B-fold. The additional signals (∼25-30) in the central region of the HSQC spectrum that are not detected in B_4_ are presumably due to the disordered C-terminal tail of S_b5_. In contrast to S_b1_, where the C-terminal tail interacts with the B-fold extensively, there appears to be less interaction in S_b5_, as evidenced by fewer changes in chemical shifts or peak intensities in the B-region of S_b5_ compared with B_4_.

### CD analysis of unfolding for S_b5_

The thermal unfolding profile of S_b5_ shows a low temperature transition with a midpoint ∼283^∘^K and a major transition with a midpoint of ∼333^∘^K (**Fig. S4D**). The NMR analysis indicates that the major transition is unfolding of the B-fold. Fitting the thermal denaturation data above 293^∘^K to the Gibbs-Helmholtz equation shows the ΔG_folding_ for the B-fold is −5kcal/mol at 298^∘^K (**Fig. 4B**). Thus the L67R mutation in S_b4_ makes the B-fold highly favorable and the S-fold highly unfavorable (>5 kcal/mol) consistent with the change in population from mixed to B-fold observed by NMR. The large shift in S-fold population between S_b4_ (∼50%) and S_b5_ (∼0%) occurs with a moderate change in the Rosetta energy for the S-fold (**Fig. 4D**), however, due to the presence of competing alternative B-states. This is discussed further below.

## Discussion

Five dimorphic sequence pairs were designed and stable structures were determined for 7 of the 10 proteins using NMR spectroscopy (**Fig. 1**). Two of the switches (one S- to A-switch and one S-to B-switch) completely achieved the goal of populating the S-fold in the longer form and the G-fold in the shorter form. We initially assumed that mutations introduced to create compatibility with two folds would necessarily compromise native state interactions in one or both of the folds. The surprising conclusion, however, is that for one S-to A-switch, and three of four S-to B-switches it was possible to design one sequence that is compatible with native state interactions for both folds. In all these cases the calculated energy of the S-fold was near the wild type sequence. This was true even though many mutations were introduced to create compatibility. It is important to understand, however, that Rosetta Relax evaluates native state interactions in the vicinity of the starting structure. The ΔG_folding_ of dimorphic proteins will also be strongly influenced by non-native states that the Relax protocol is not evaluating.

Engineering stability of an antagonistic, embedded fold necessarily destabilizes the longer fold even when native-state interactions are not compromised. Consequently, three basic structural transitions dictate the behavior of a dimorphic protein. To understand these transitions, it is useful to divide the structure of the longer protein into the sequence corresponding to the 56 residue embedded fold (part 1) and the remaining sequence (part 2). Both parts are ordered in the S-fold. When part 1 switches into a G-fold, however, part 2 unfolds. The conformations of part 1 are denoted s1 (S-conformation), g1 (A- or B-conformation), and u1 (unfolded conformation). The conformations of part 2 are denoted s2 (S-conformation) and u2 (unfolded conformation). Consider the equilibria:

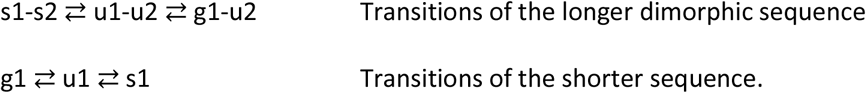

In an idealized dimorphic protein, the energy from native interactions in both S- and G-folds would be equivalent to native interactions in the two natural proteins. If we then assume that part 2 only interacts with s1 to form s1-s2 or with solvent in g1-u2 and u1-u2 forms, then we can predict the populations of all the species in the switch from the equilibrium constants for the S-fold (K_S_) and the A-fold (K_A_). These are calculated using the ΔG_folding_ of the wild type proteins (**Fig. 4**) and the Gibbs equation ΔG = -RT ln (K). For example, the expected populations in an idealized switch from the S-to the A-fold would be:

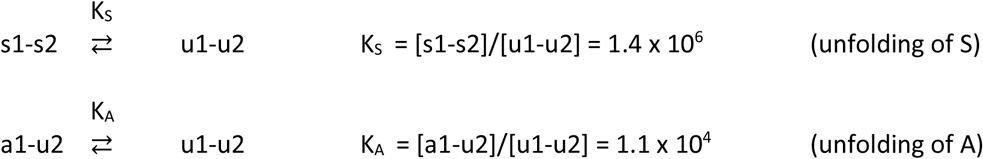

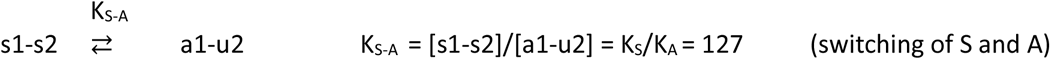

Rules for switches emerge from examining deviations from idealized behavior. Each of the five dimorphic sequences were assessed for how well the experimental structure matches the design and how well the switching energetics match the idealized case. We observe behaviors ranging from near ideal to large deviations from ideal, but surprisingly, most deviations do not appear to result from compromised native interactions in S- and G-folds. Rather, deviations appear to arise from promiscuous interactions of u2 with alternative folds of part 1. That is, the assumption that u2 interacts only with s1 to form an S-fold or with solvent in an unfolded state is invalid. In fact, s2 forms alternative interactions with the G-fold that compete with formation of the S-fold.

### Case 1, S_a1_ to A_1_ switch

Part 1 in this switch comprises residues 11-66 and part 2 residues 1-10 and 67-95. The experimental and designed structures of S_a1_ match (**Fig. 2**), Rosetta energies are similar, and the observed ΔG_folding_ of S_a1_ is −7.5 kcal/mol, compared to an idealized value of −8.5 kcal/mol. Likewise, the 56 amino acid A_1_ protein has a stable structure that closely matches the designed model. This example comes closest to an idealized case. This switching behavior occurs because the S-fold is considerably more stable than the embedded, antagonistic A-fold and the equilibrium strongly favors S in the longer protein (**Fig. 4C**).

### Case 2, S_b1_ to B_1_ switch

Part 1 in this switch comprises residues 5-60 and part 2 residues 1-4 and 61-95. The designed and experimental structures of B_1_ match quite well in this case, but those of S_b1_ do not (**Fig. 5**). In fact, S_b1_ populates a B-like fold even though the Rosetta energy of the S_b1_ design model is only a little less than native S protein (**Fig. 4D**). Close examination of the ensemble of NMR structures for S_b1_, shows that part 2 of the protein has propensity for S-type structures even when part 1 has a B-fold. Thus, the observed conformational ensemble can be denoted g1-s2. This ensemble is populated because stability of the internal g1-state compromises the stability of the s1-s2-fold and results in promiscuous interactions of s2 with g1. The overall ΔG_folding_ for the ensemble of B-like folds in S_b1_ is −1.1 kcal/mol. Since no S-fold is observed in S_b1_, this means that the ΔG_folding_ for the S-fold is >1.1 kcal/mol (**Fig. 4C**).

### Case 3, S_b2_ to B_2_ switch

To generate a second version of this switch, we introduced 13 mutations into the S_b1_ sequence. The mutations result in a switch from a B-fold into an S-fold and the designed and experimental structures of S_b2_ roughly match (**Fig. S6**). The observed ΔG_folding_ of S_b2_ is −4.0 kcal/mol. The switch from B- to S-folds does not appear to arise from improved native interactions in the S-fold, however. In fact, the Rosetta energy of the S_b2_ design model is almost identical to the S_b1_ design model (**Fig. 4D**). Rather, the B-to S-switch results from decreased stability of the antagonistic, embedded B-fold in S_b2_.

### Case 4, S_b3_ to B_3_ switch

Part 1 in this switch comprises residues 1-56 and part 2 residues 57-87. The designed and experimental structures roughly match (**Fig. 6**). S_b3_ populates an S-fold, although deviations exist in loops. The Rosetta energy of the S_b3_ design model is very similar to that of the natural sequence (**Fig. 4D**). The observed ΔG_folding_ of the S-fold is only −3.5 kcal/mol, however, and shows the influence of the antagonistic B-fold on S-fold stability. B_3_ primarily populates a B-fold, but similarly its low stability (−1.2 kcal/mol) may indicate some propensity for the antagonistic S-conformation.

### Case 5 S_b4_ and S_b5_ to B_4_ switch

The Y5L mutation introduced into S_b3_ results in simultaneous population of multiple, folded conformational states. This appears to arise from slightly compromised native interactions in the S-fold and increased stability of the antagonistic, embedded B-fold. The 56 amino acid B_4_ protein has a B-fold that matches the designed model. The ΔG_folding_ of B_4_ is −4.1 kcal/mol compared to −6.6 kcal/mol for the natural G_B_ protein. This is an example in which good computational design produces a complex result. The complexity appears to arise because the stability of the embedded B-fold and the longer S-folds are similar and antagonistic. This causes more than one fold to be populated and the observed stability of the S-fold to be lower than predicted based on its *REU* value. S_b4_ is thus at a critical point in switching between the S- and B-folds. Consequently, single substitution mutations in S_b4_ can produce either a stable S-fold or a stable B-fold (**Fig. 8**).

**Figure 8:**
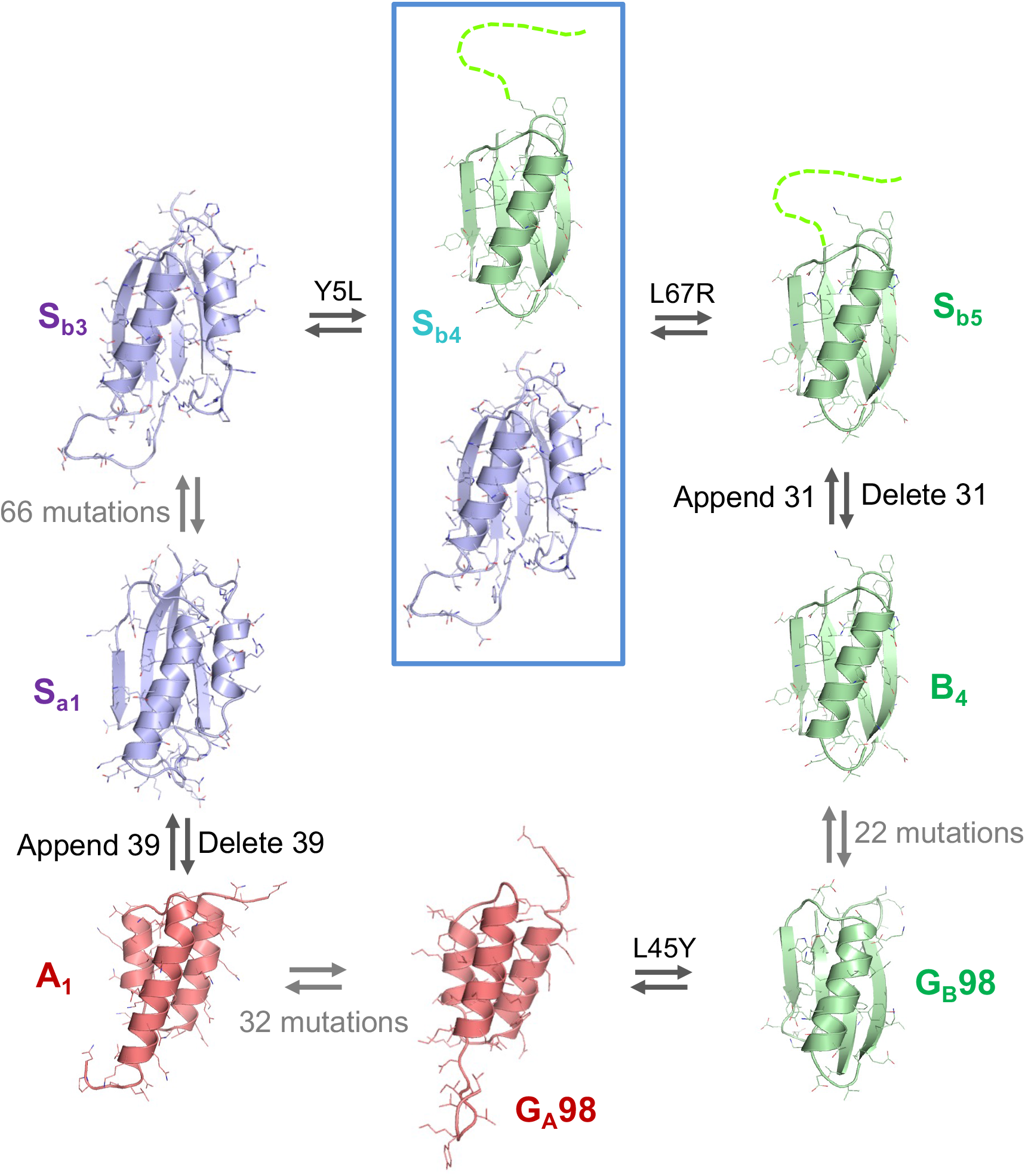
Sequence-fold relationships of engineered S-, A-, and B-folds. Switches between stable folds can be induced by point mutation or deleting/appending the part 2 sequence. Blue denotes an S-fold, green a B-fold, and red an A-fold. Gray arrows connect proteins that have been reengineered without a fold switch. S_b4_ (blue box) populates two folds simultaneously. The G_A_98 and G_B_98 proteins were described in an earlier paper (8).

Several general rules for engineering switches emerge from this study. First, it is possible to design one sequence with good native state interactions in two folds. If this condition is met, then the two main factors determining fold populations are the stability of the embedded G-fold relative to the S-fold and the conformational propensities of the ends that are generated in the switch to the embedded fold. The higher the stability of the embedded fold relative to the larger fold, the more the ends are populated, and the more the structures deviate from design (e.g. S_b1_). Thus, the ends generated in a switch are a double-edged sword. They are a repository of switching energy to drive the G-fold to the S-fold but also can contribute energy to switch to other states. Finally, successful design of a dimorphic sequence creates a critical state in which conformation is extremely sensitive to small perturbations anywhere in the sequence. Thus, as in other complex systems, a small change may have a “butterfly effect” on how the folds are populated (**Fig. 8**).

Rules for engineering switches also highlight several well-established principles of protein folding. First, the stability of the non-native state, though less predictable, contributes to overall stability as much as the native state (53-60). Second, the effect of the appended ends is often denaturing. Ends can be generally denaturing by forming non-specific backbone hydrogen bonds and hydrophobic interactions in the unfolded state ensemble, but promiscuous interactions of structured end fragments may also result in alternative structures (61). This study also shows a surprising attribute of the folding code: It is not difficult to engineer sequences that are compatible with native interactions in more than one fold. This is consistent with the existence of natural dimorphic proteins, but the fact remains that most natural protein sequences populate only one fold (62). We suggest that evolutionary pressure to avoid critical states typically causes proteins to evolve toward a single native state. Thus, two-state behavior may result from nature’s negative design rather than being an inherent property of the protein code (63-65). Proteins generally evolve toward two-state behavior because dimorphic sequences can generate promiscuous interactions that destabilize them, compromise their function, and may be pathological.

## Materials and Methods

### Mutagenesis, protein expression and purification

Mutagenesis was carried out using Q5^®^ Site-Directed Mutagenesis Kits (NEB). G_A_ and G_B_ variants were cloned into a vector (pA-YRGL) encoding the sequence:

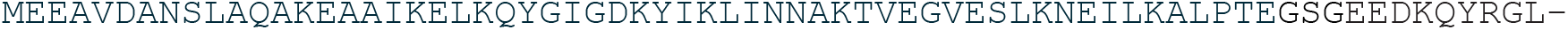

as an N-terminal fusion domain (39). The resulting fusion proteins were purified using a second generation of the affinity-cleavage tag system used previous to purify switch proteins (8, 66). The second generation tag (*YRGL-tag*) results in high-level soluble expression of the switch proteins and also enables capture of the fusion protein by binding tightly to an immobilized processing protease via the C-terminal EEDQYRGL sequence. The addition of 100µM imidazole activates an immobilized, imidazole-activated protease (*Im-Prot*) and releases the purified switch protein from the *Im-Prot* media (Potomac Affinity Proteins). The purified protein was then concentrated to 0.2 to 0.3 mM, as required for NMR analysis. The purification system is described in detail in Supplemental methods and is available from Potomac Affinity Proteins.

### Circular Dichroism (CD)

CD measurements were performed in 100mM KPO_4_, pH 7.2 with a Jasco spectropolarimeter, model J-1100 with a Peltier temperature controller. Quartz cells with path lengths of 0.1 cm and 1cm were used for protein concentrations of 3 and 30 µM, respectively. The ellipticity results were expressed as mean residue ellipticity, [θ], deg cm2 dmol-1. Ellipticities at 222 nm were continuously monitored at a scanning rate 0.5 deg/min. Reversibility of denaturation was confirmed by comparing the CD spectra at 293^∘^K before melting and after heating to 373^∘^K and cooling to 293^∘^K.

### NMR Spectroscopy

Isotope-labeled samples were prepared at 0.2-0.3 mM concentrations in 100 mM potassium phosphate buffer (pH 7.0) containing 5% D_2_O. NMR spectra were collected on Bruker AVANCE III 600 and 900 MHz spectrometers fitted with Z-gradient ^1^H/^13^C/^15^N triple resonance cryoprobes. Standard double and triple resonance experiments (HNCACB, CBCA(CO)NH, HNCO, HN(CA)CO, and HNHA) were utilized to determine main chain NMR assignments. Inter-proton distances were obtained from 3D ^15^N-edited NOESY and 3D ^13^C-edited NOESY spectra with a mixing time of 150 ms. NmrPipe (67) was used for data processing and analysis was done with Sparky (68). Two-dimensional {^1^H}-^15^N steady state heteronuclear NOE experiments were acquired with a 5 s relaxation delay between experiments. Chemical shift perturbations between B_1_ and S_b1_ were calculated using Δδ_total_=((*W*_H_Δδ_H_)^2^ + (*W*_N_ΔδN)^2^)^1/2^, where Δδ_H_ and Δδ_N_ represent ^1^H and ^15^N chemical shift changes, respectively. For PRE experiments on S_b1_, single-site cysteine mutant samples were incubated with 10 equivalents of MTSL ((1-oxyl-2,2,5,5-tetramethylpyrroline-3-methyl) methanethiosulfonate, Santa Cruz Biotechnology) at 25°C for 1 hour and completion of labeling was confirmed by MALDI mass spectrometry. Control samples were reduced with 10 equivalents of sodium ascorbate. Backbone amide peak intensities of the oxidized and reduced states were analyzed using Sparky. Three-dimensional structures were calculated with CS-Rosetta3.2 using experimental backbone ^15^N, ^1^H_N_, ^1^Hα ^13^Cα, ^13^Cβ, and ^13^CO chemical shift restraints and were either validated by comparison with experimental backbone NOE patterns (A_1_, B_1_, B_4_, S_b1_) or directly employed interproton NOEs (S_a1_, S_b2_) or PREs (S_b1_) as additional restraints. One thousand CS-Rosetta structures were calculated from which the 10 lowest energy structures were chosen. For S_b3_, CS-Rosetta failed to converge to a unique low energy topology, producing an approximately even mixture of S- and B-type folds despite the chemical shifts and NOE pattern indicating an S-fold. In this case, CNS1.1 (69) was employed to determine the structure as described previously (39), including backbone dihedral restraints from chemical shift data using TALOS (70). Protein structures were displayed and analyzed utilizing PROCHECK-NMR (71), MOLMOL (72) and PyMol (Schrodinger) (38).

## Supporting information

Supporting data, analysis, and methods

## Acknowledgments

This work was supported by National Institutes of Health Grant GM62154 (to PB and JO) and 5R44GM126676 (to PB). The NMR facility is supported by the University of Maryland, the National Institute of Standards and Technology, and a grant from the W. M. Keck Foundation. We would also like to thank Dr. Nese Sari for her thoughtful comments. Mention of commercial products does not imply recommendation or endorsement by NIST.

