## Supporting data, analysis, and methods for "Rules for designing protein fold switches and their implications for the folding code"

**Supplemental Material**

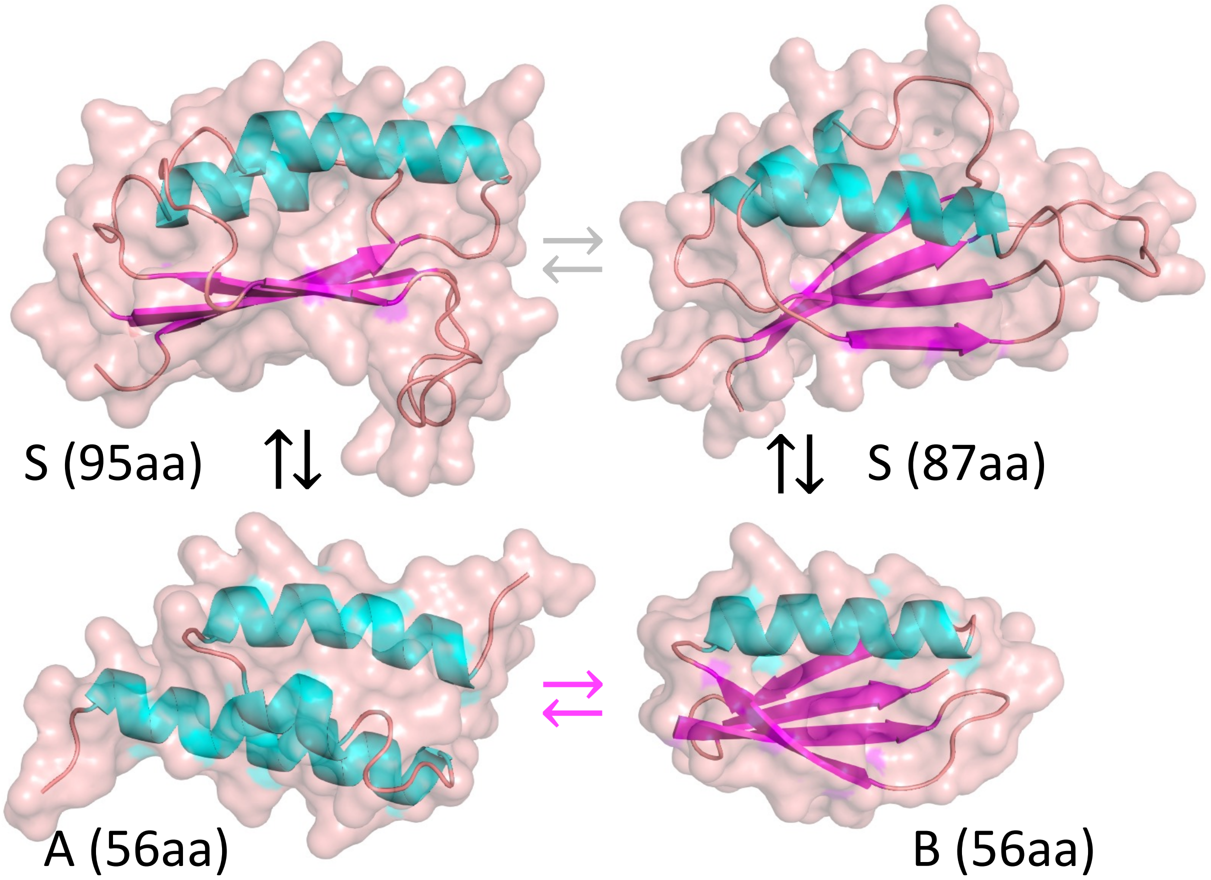


| parent protein | Engineered protein | Fold type | aa | Natural function |
|---|---|---|---|---|
| $G_A$ domain of Protein G | $A_1$ | ααα | 56 | Albumin binding |
| $G_B$ domain of Protein G | $B_1$, $B_2$, $B_3$, $B_4$ | ββαββ | 56 | IgG binding |
| S6 | $S_{a1}$, $S_{b1}$, $S_{b2}$ | βαββαβ | 95 | RNA binding |
| S (loop variant) | $S_{b3}$, $S_{b4}$, $S_{b5}$ | βαββαβ | 87 | Protease inhibitor |

**Figure S1.** Switching among three common folds. Black arrows connect structures of proteins with dual propensity for S- and A-folds and S- and B-folds. The magenta arrows indicate that a dual propensity sequence has previously been engineered connecting the A- and B-folds (8). The grey arrows indicate the connection between the native S6 protein and a loop variant in the S- superfold family (10, 50).

### $A_1$ / $S_{a1}$ switch

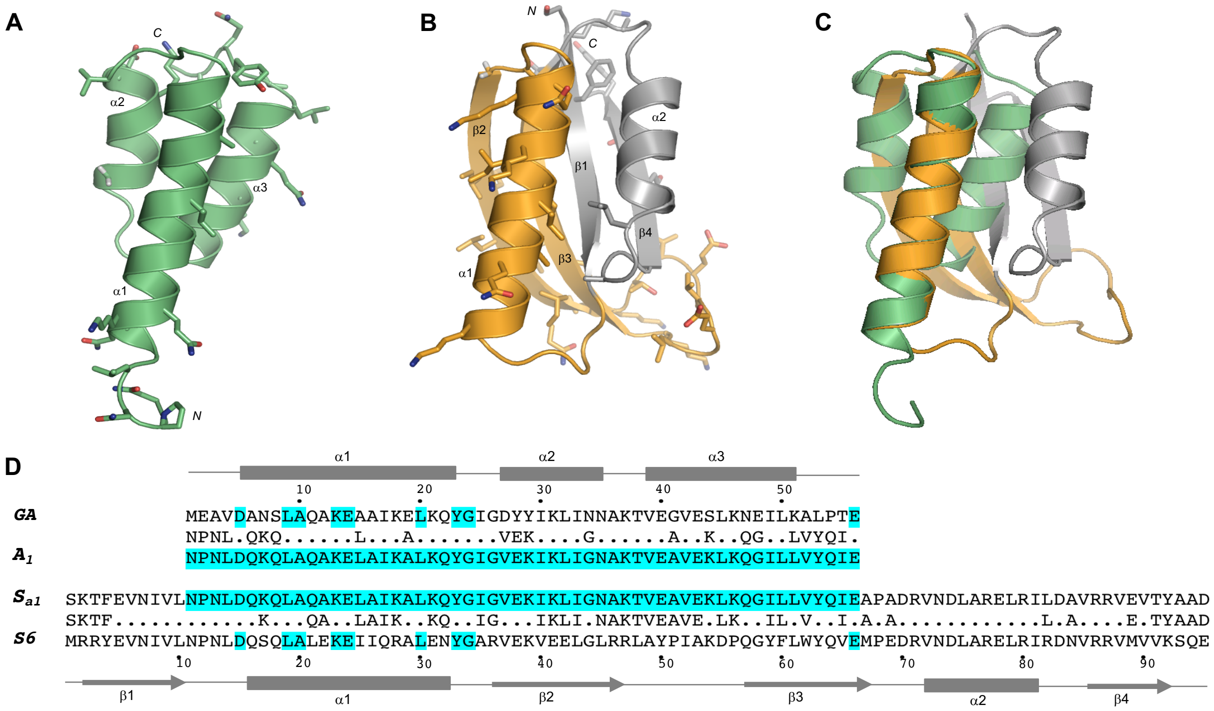


**Figure S2:** Summary of mutations made in $G_A$ and S6 parent folds to achieve high identity. (**A**) Mutations made in the $G_A$ parent structure (PDB 2FS1) (1). (**B**) Mutations made in the S6 parent structure (PDB 1RIS). The region co-evolved to high sequence identity with $G_A$ is highlighted (orange). (**C**) Structural alignment using the α1-helices of $G_A$ and S6. (**D**) Corresponding sequence alignment. Residues highlighted in blue in the parent sequences indicate original positions of identity between $G_A$ and S6. Residues highlighted in blue in the designed sequences indicate identity between $A_1$ and $S_{a1}$. The secondary structures of the parent $G_A$ and S6 sequences are shown above and below the alignments, respectively.

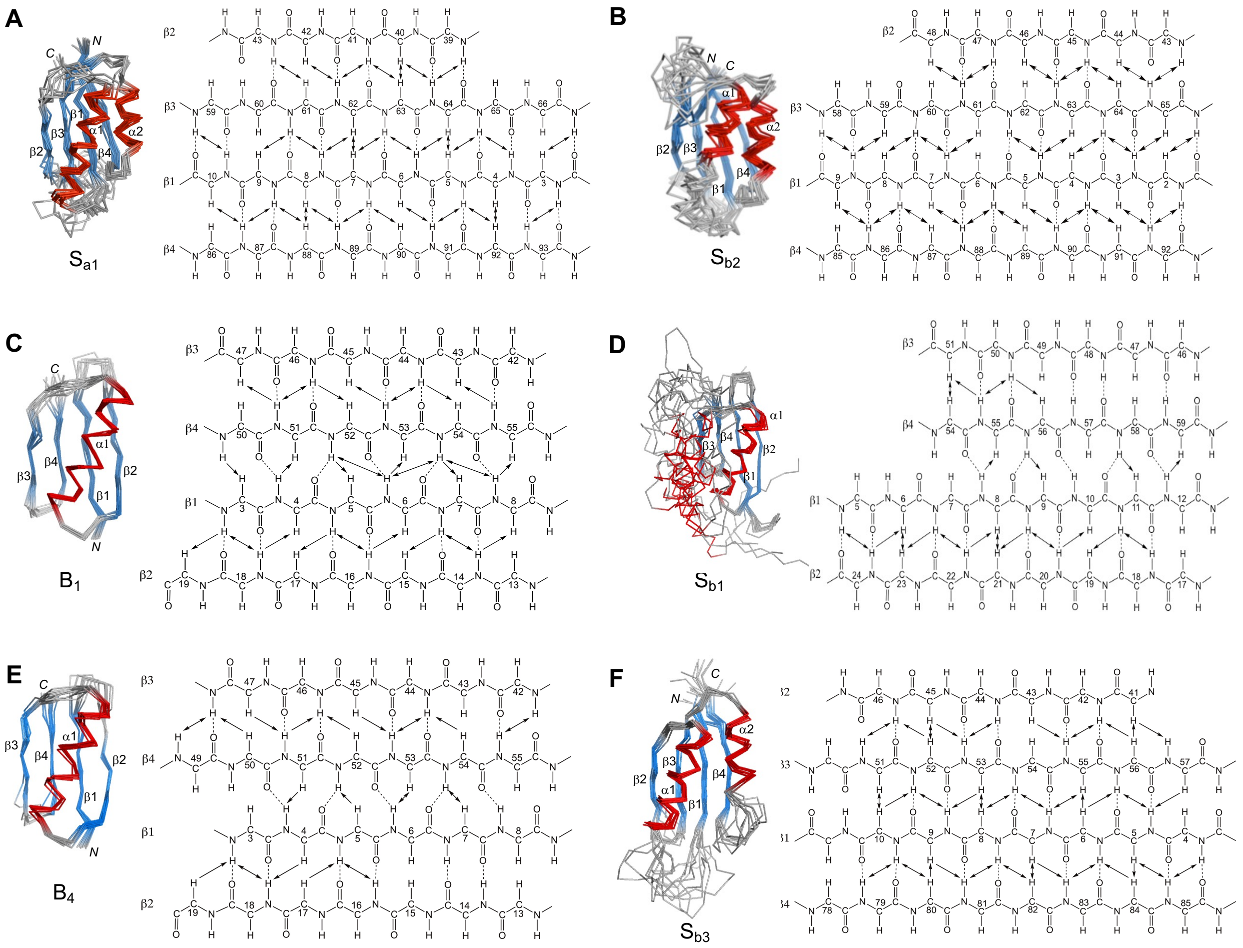


**Figure S3**: Summary of long-range backbone NOEs observed for β-sheets in designed proteins.

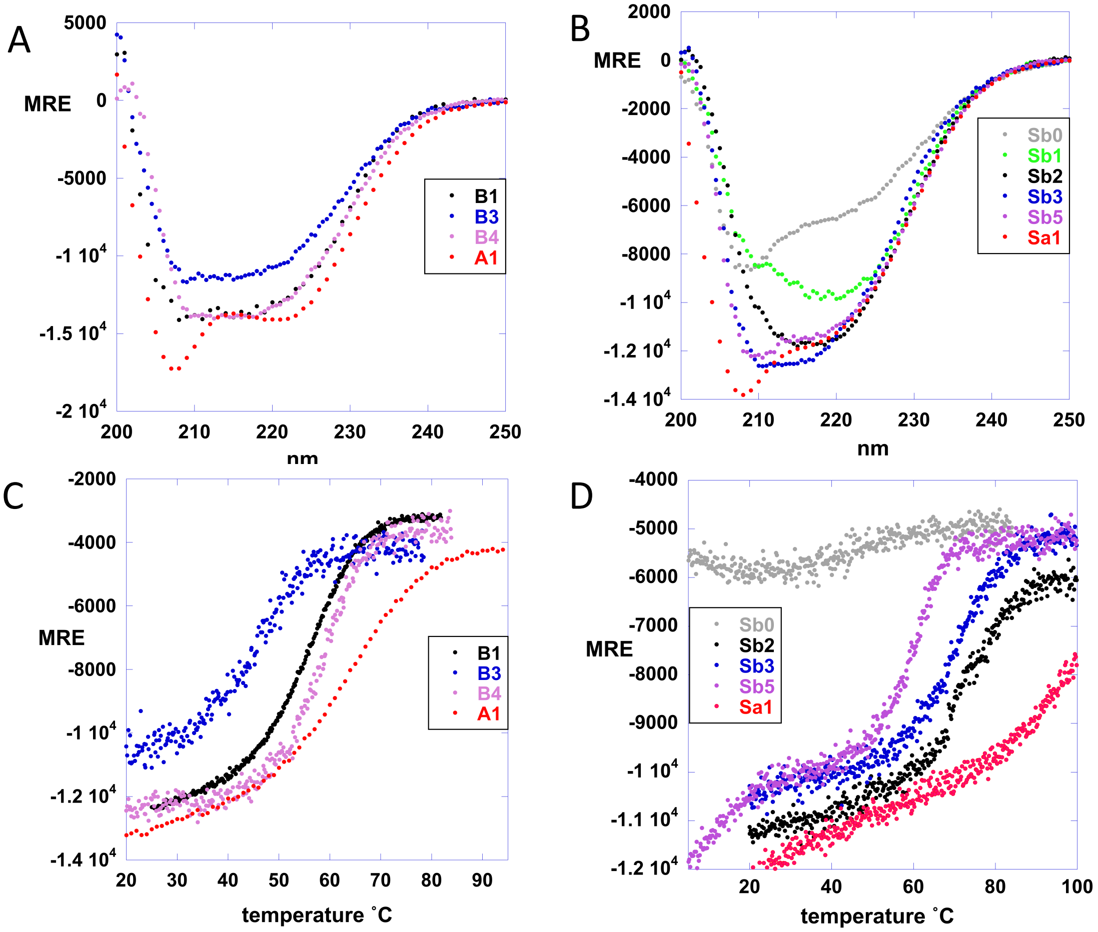


**Figure S4:** CD spectra/melting curves. (**A**) CD spectra for 56 amino acid proteins. (**B**) CD spectra for 87 and 95 amino acid proteins. (**C**) Ellipticity at 222nm plot vs. temperature for 56 amino acid proteins. (**D**) Ellipticity at 222nm plot vs. temperature for 87 and 95 amino acid proteins. $S_{b0}$ is an unstable variant (F7V) of $S_{b1}$ used to measure the temperature dependence of the unfolded state.

### $B_1$ / $S_{b1}$ and $B_1$ / $S_{b2}$ switch (80% sequence identity)

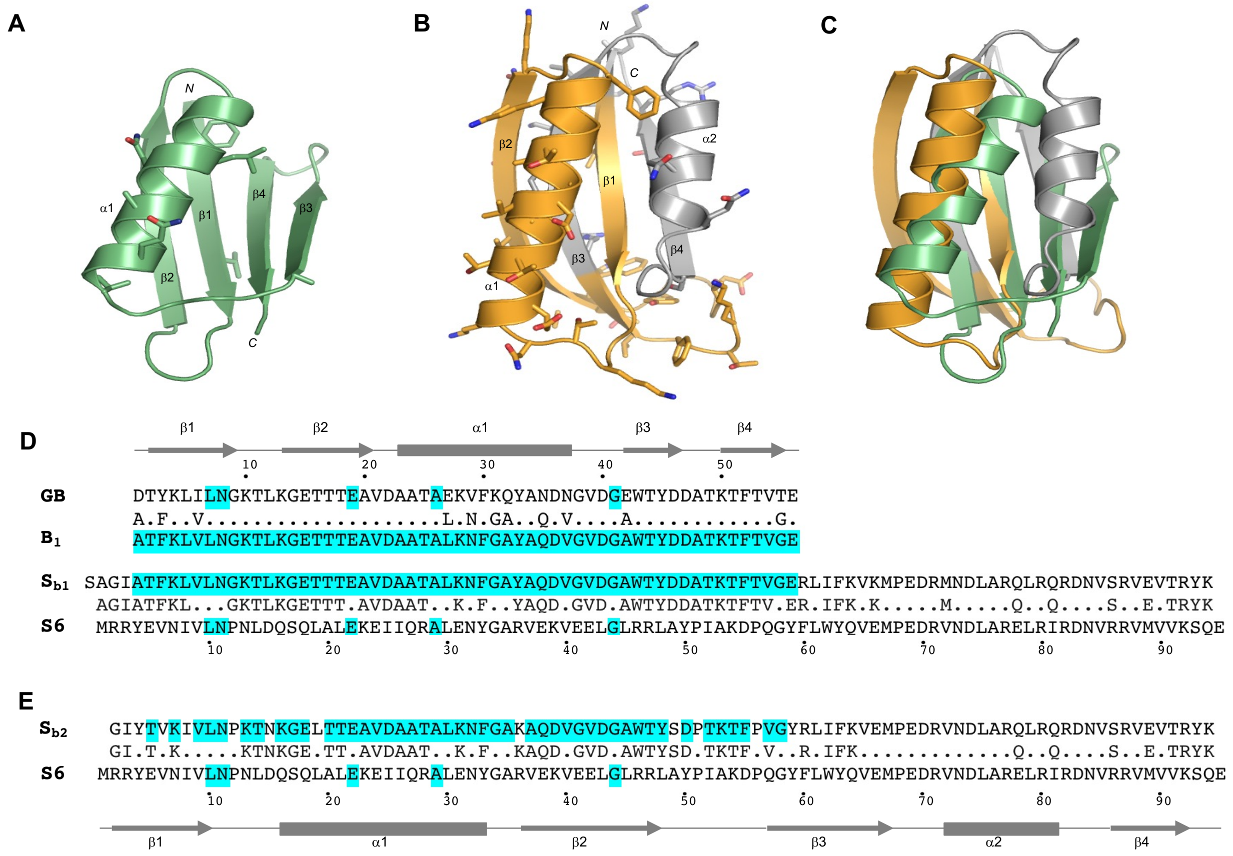


**Figure S5**: Summary of mutations made in $G_B$ and S6 parent folds to achieve 80% sequence identity. (**A**) Residues mutated in the $G_B$ parent structure (PDB 1PGA) are highlighted. (**B**) Residues mutated in the S6 parent structure (PDB 1RIS). The region co-evolved to high sequence identity with $G_B$ is highlighted (orange). (**C**) Structural alignment using the β1–strands of $G_B$ and S6. (**D**) Sequence alignment of $B_1$ and $S_{b1}$. Residues highlighted in blue in the parent sequences indicate positions of identity between $G_B$ and S6. $B_1$ and $S_{b1}$ are identical over the entire $B_1$ sequence. (**E**) Sequence alignment of $S_{b2}$ with the parent S6. Residues highlighted in $S_{b2}$ indicate identity between $S_{b2}$ and $B_1$, which is 80%. The secondary structures of the parent $G_B$ and S6 sequences are shown above and below the alignments, respectively.

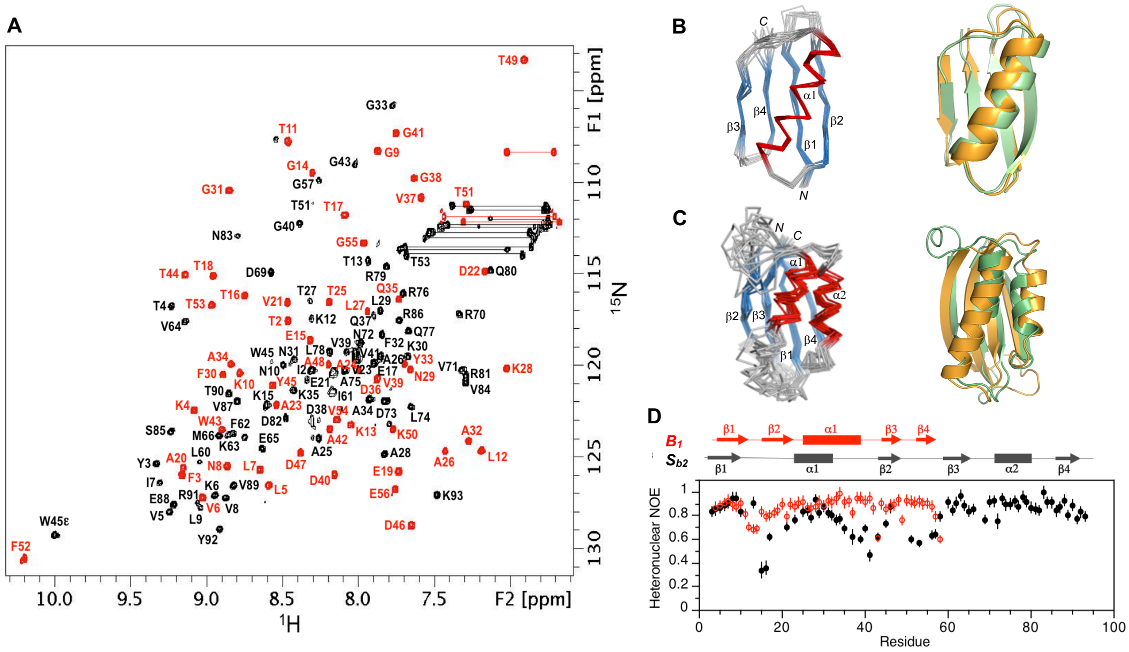


**Figure S6**: Structure and dynamics of $B_1$ and $S_{b2}$. (**A**) Overlaid two dimensional $^1H$-$^{15}N$ HSQC spectra of $S_{b2}$ (black) and $B_1$ (red) with backbone amide assignments. Spectra were recorded at 298°K. (**B**) Ensemble of 10 lowest energy CS-Rosetta structures for $B_1$ (left panel). Superposition of the $B_1$ structure (green) with the parent $G_B$ fold (orange) (right panel). (**C**) Ensemble of 10 lowest energy CS-Rosetta structures for $S_{b2}$ (left panel). Superposition of $S_{b2}$ (green) with the parent S6 fold (orange) (right panel). (**D**) Plot of {$^1H$}-$^{15}N$ steady state heteronuclear NOE values at 600 MHz versus residue for $B_1$ (red) and for $S_{b2}$ (black). Error bars indicate ±1SD.

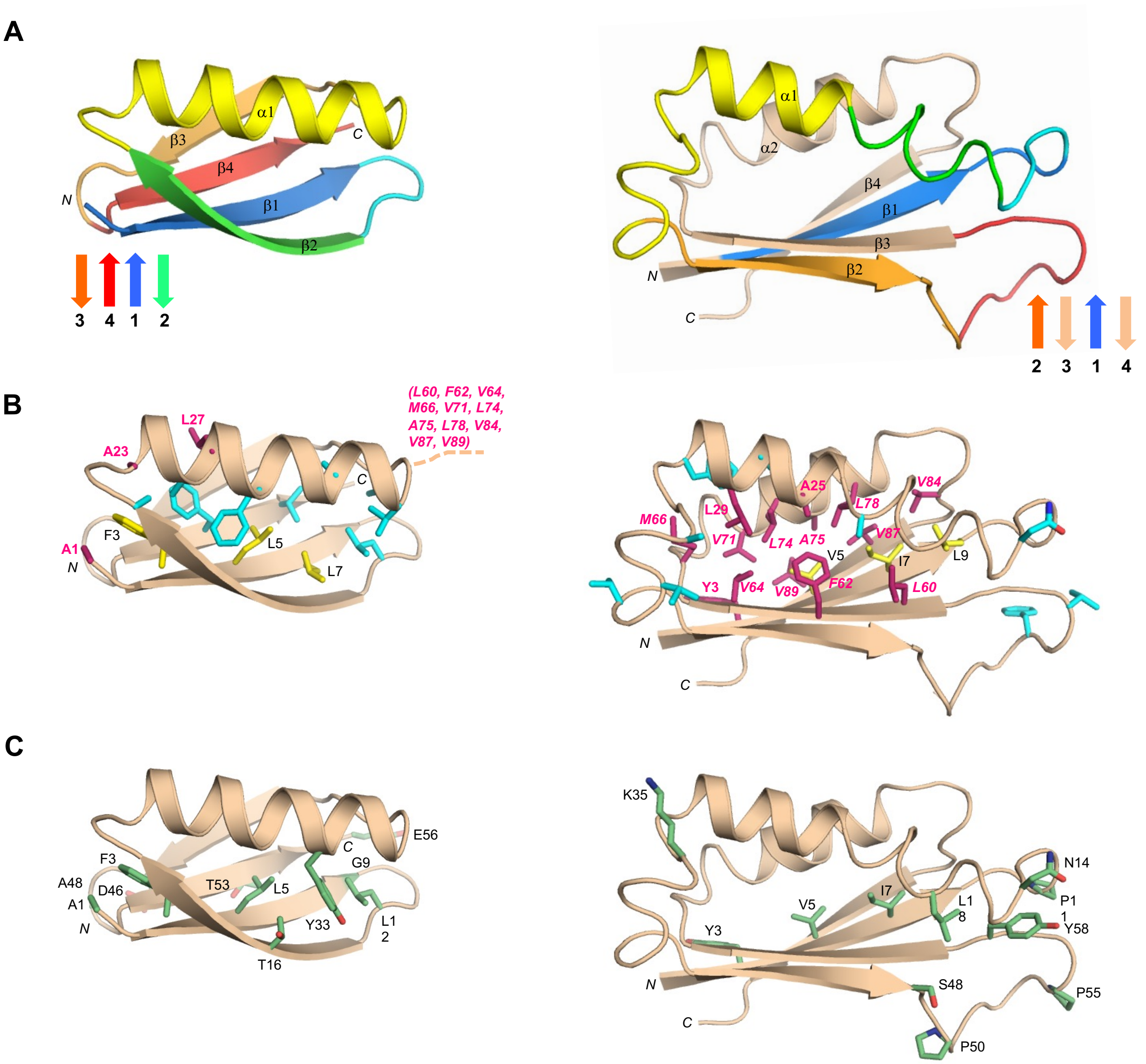


**Figure S7**: Structural differences in the high sequence identity regions of $B_1$ and $S_{b2}$. (**A**) Main chain comparisons. (Left panel) CS-Rosetta structure of $B_1$ with color coding for regions containing secondary structured elements. (Right panel) Corresponding color coded regions mapped onto the CS-Rosetta structure of $S_{b2}$, illustrating changes in backbone conformation. Regions outside the 56 amino acid sequence of $B_1$ are shown in wheat. Topology diagrams with the arrangement of β-strands are shown next to each structure. (**B**) Side chain comparisons. (Left panel) Residues contributing to the core of $B_1$ from the β1-strand (yellow), and from other regions (cyan). The non-β1 core residues from $S_{b2}$ (pink) do not overlap with the $B_1$ core (see text for further details). (Right panel) Residues contributing to the core of $S_{b2}$ from the β1-strand (yellow), and most of the other participating core residues (pink). The non-β1 core residues from $B_1$ are also shown (cyan), highlighting the low degree of overlap. (**C**) Positions of amino acid differences between $B_1$ and $S_{b2}$ in the overlapping 56 amino acid region.

### $B_4$ / $S_{b3}$ switch (98% sequence identity)

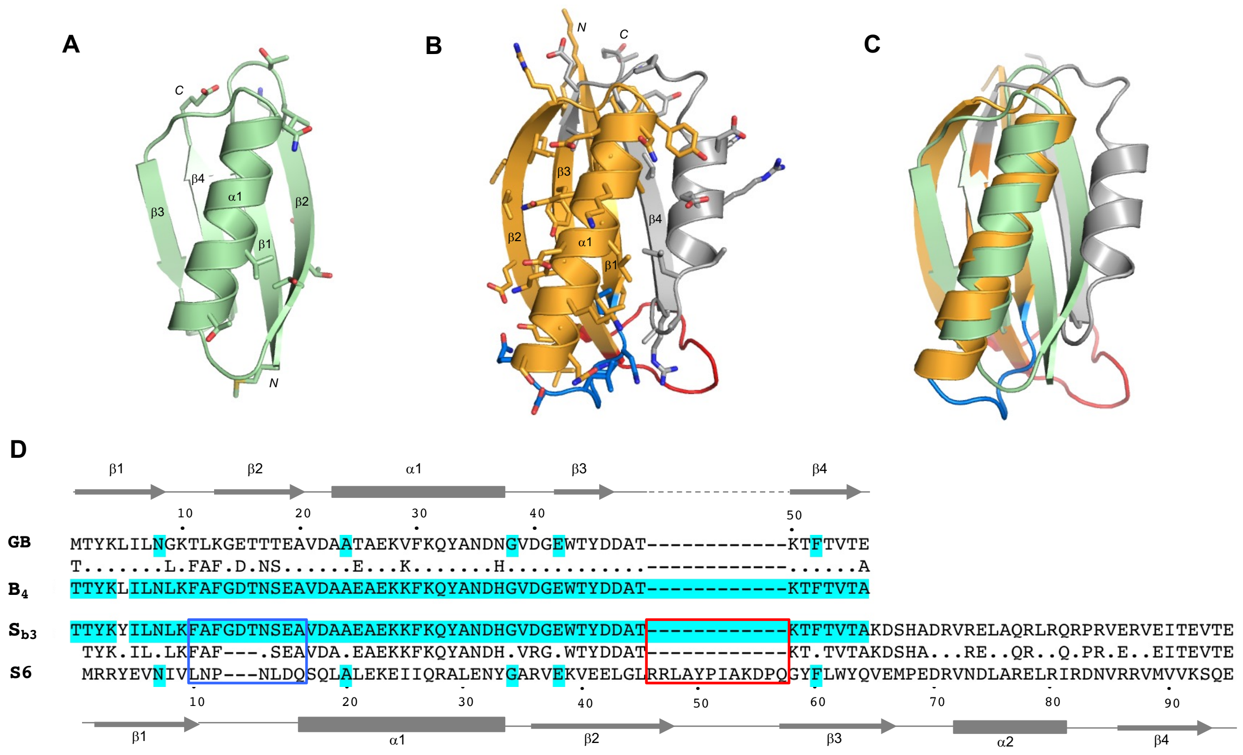


**Figure S8**: Summary of mutations made in $G_B$ and S6 parent folds to achieve 98% sequence identity. (**A**) Mutations made in the $G_B$ parent structure (PDB 1PGA). (**B**) Mutations made in the S6 parent structure. The region co-evolved to high sequence identity with $G_B$ is highlighted (orange). Insertion and deletion loops are shown in blue and red, respectively. **(C)** Structural alignment using the $\alpha1/\beta3/\beta4$ and $\alpha1/\beta2/\beta3$ regions of the $G_B$ and S6 folds, respectively. (**D**) Corresponding sequence alignment. Residues highlighted in blue in the parent sequences, $G_B$ and S6, indicate positions of initial identity. Residues highlighted in blue in the designed sequences indicate the final level of identity between $B_4$ and $S_{b3}$. The positions where residues were inserted and deleted are shown in blue and red boxes, respectively. The secondary structures of the parent $G_B$ and S6 sequences are shown above and below the alignments, respectively.

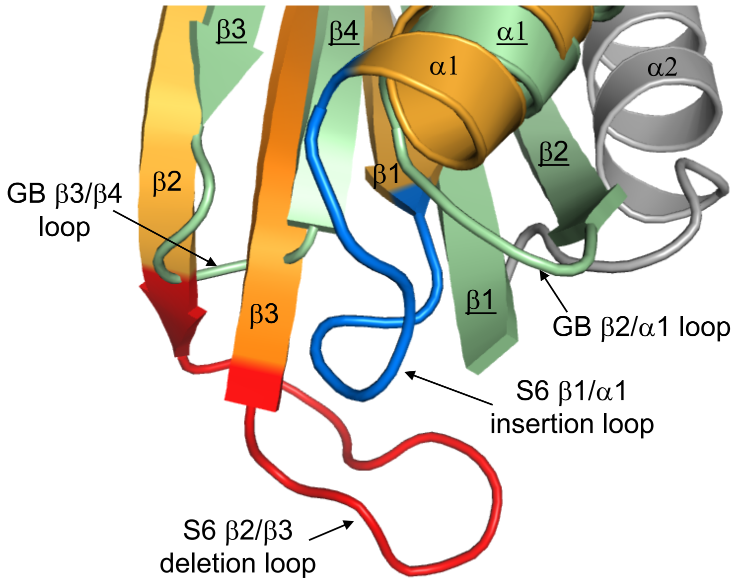


**Figure S9**: Superposition of $G_B$ (green) and S6 (orange/gray) folds highlighting the S6 deletion (red) and insertion (blue) loops.

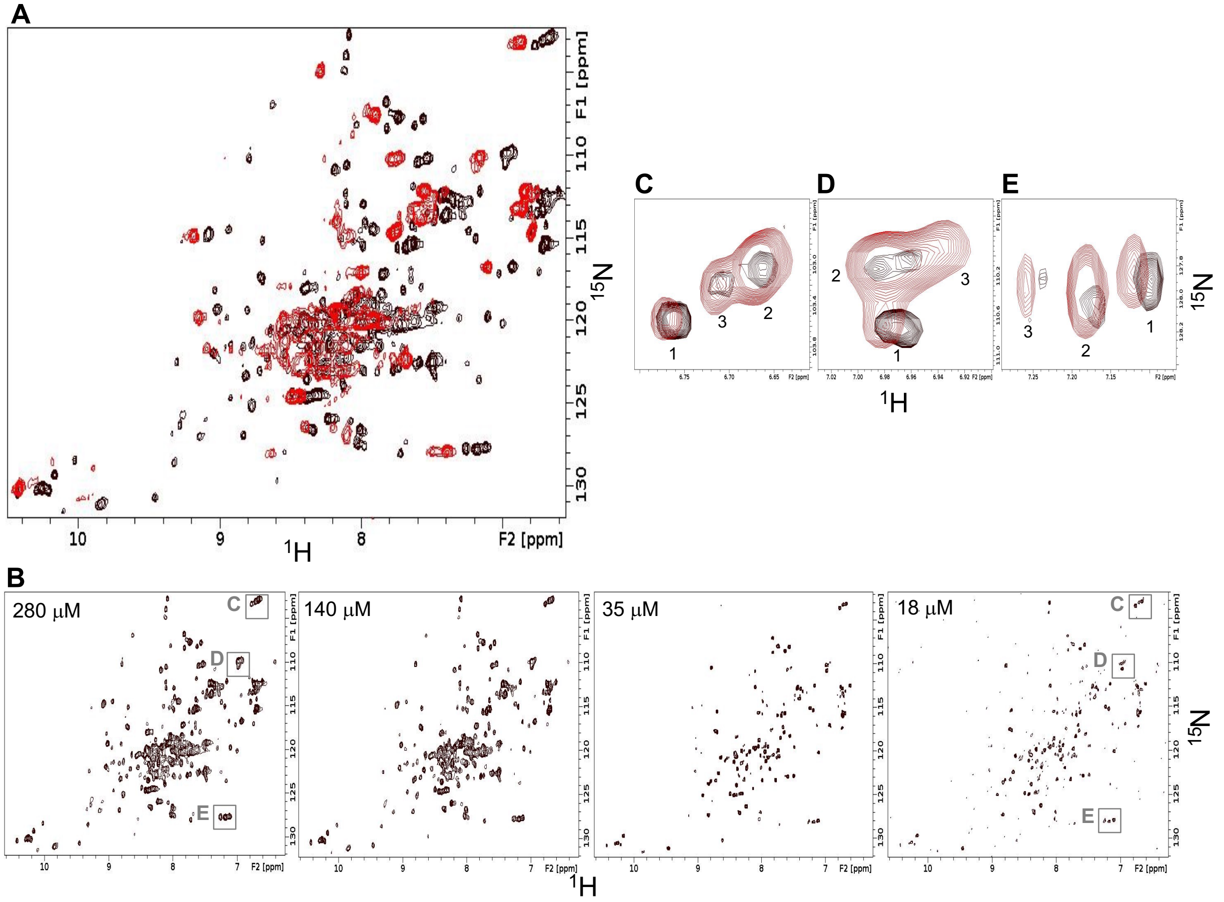


**Figure S10:** Temperature and concentration dependence of $B_3$. (**A**) Overlaid two dimensional $^1$H-$^{15}$N HSQC spectra of $B_3$ at 298$^o$K (red) and at 278$^o$K (black). The protein concentration in both spectra is 280 μM. Broadened peaks at 298$^o$K are consistent with low stability. (**B**) HSQC spectra of $B_3$ recorded at different concentrations as indicated. All spectra were acquired at 278$^o$K. (**C, D, E**) Expanded regions from the overlaid concentration dependent spectra of $B_3$ at 18 μM (black) and 280 μM (red). Regions are as indicated in the leftmost and rightmost panels in (B). The peaks in (C, D, E) are labeled 1, 2 and 3, corresponding to three main concentration-dependent species. At 18 μM, the monomeric species 1 is dominant, while at 280 μM the putatively dimeric species 2 is dominant.

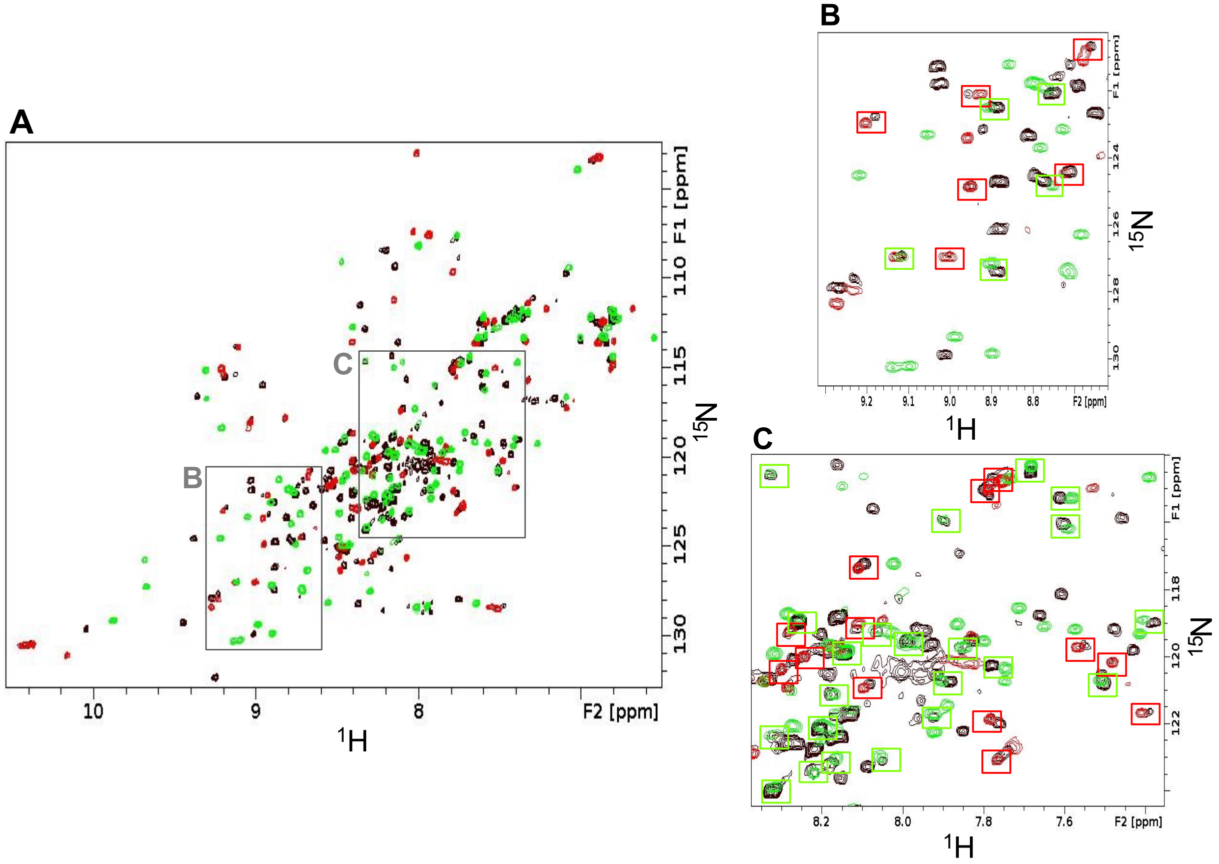


**Figure S11:** Comparison of spectra for $S_{b4}$, $B_4$, and $S_{b3}$. **(A)** Two dimensional $^{1}H$-$^{15}N$ HSQC spectrum of $S_{b4}$ (black) overlaid with spectra for $B_4$ (red) and $S_{b3}$ (green). All spectra were acquired at 298°K. (**B, C**) Expanded regions as indicated in (**A**). Overlapping or proximal $B_4$/$S_{b4}$ peaks (red boxes) and $S_{b3}$/$S_{b4}$ peaks (green boxes) are highlighted. The overlap of the spectra indicates that $S_{b4}$ populates both the S- and B-folds simultaneously. About 50 peaks in $S_{b4}$ superimpose well with $B_4$ although some peaks are shifted slightly, presumably due to the presence of a disordered C-terminal tail in $S_{b4}$ when the polypeptide chain adopts the B-fold. A significant number of $S_{b4}$ signals also coincide approximately with the $S_{b3}$ spectrum. Differences in peak positions between $S_{b4}$ and $S_{b3}$ are likely due to the L5Y mutation, which affects numerous neighboring residues.

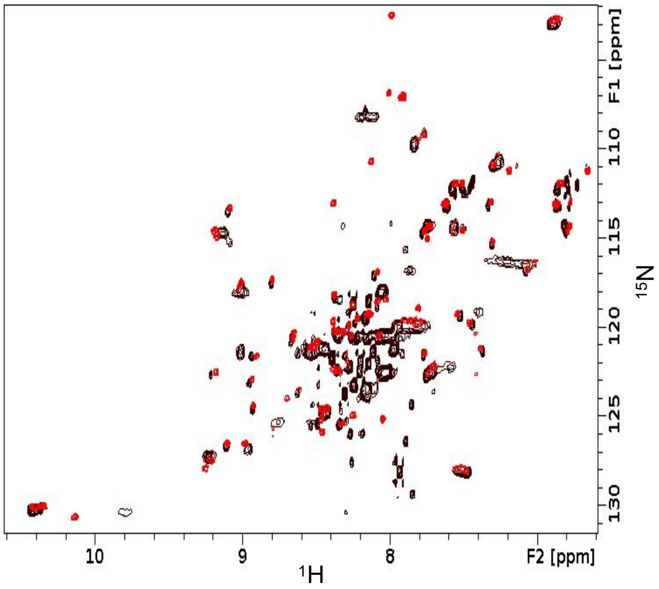


**Figure S12**: Two dimensional $^{1}H$-$^{15}N$ HSQC spectrum of $S_{b5}$ (black) compared with $B_4$ (red). Both spectra were acquired at 298°K. The significant overlap of folded signals in the two spectra indicates that $S_{b5}$ has a B-fold with a disordered C-terminal tail (see text for more details).

**Table S1**: Structure statistics for $A_1$, $B_1$, and $B_4$.

| | $A_1$ | $B_1$ | $B_4$ |
|---|---|---|---|
| *A. Experimental chemical shift inputs* | | | |
| $^{13}C\alpha$ | 55 | 56 | 56 |
| $^{13}C\beta$ | 51 | 50 | 53 |
| $^{13}CO$ | 54 | 55 | 53 |
| $^{15}N$ | 54 | 55 | 54 |
| $^{1}H_N$ | 54 | 55 | 54 |
| $^{1}H\alpha$ | - | 55 | - |
| *B. RMSDs to the mean structure (Å)* | | | |
| Over all residues | | | |
| Backbone atoms | 1.73±0.47 | 0.86±0.22 | 0.85±0.34 |
| Heavy atoms | 2.27±0.53 | 1.37±0.29 | 1.43±0.45 |
| Secondary structures[a] | | | |
| Backbone atoms | 0.90±0.29 | 0.67±0.22 | 0.62±0.25 |
| Heavy atoms | 1.55±0.42 | 1.12±0.25 | 1.33±0.41 |
| *C. Measures of structure quality (%)* | | | |
| Ramachandran distribution | | | |
| Most favored | 98.78±1.64 | 92.31±3.35 | 92.36±2.71 |
| Additionally allowed | 1.22±1.64 | 7.69±3.36 | 7.64±2.71 |
| Generously allowed | 0.00±0.00 | 0.00±0.00 | 0.00±0.00 |
| Disallowed | 0.00±0.00 | 0.00±0.00 | 0.00±0.00 |
| *D. Backbone RMSDs to the parent structure[b] (Å)* | | | |
| Over all residues | 2.48 | 0.62 | 0.64 |
| Secondary structures | 1.21 | 0.57 | 0.49 |
| *E. PDB/BMRB codes* | | | |
| PDBDEV | 00000083 | 00000084 | 00000085 |
| BMRB | 50907 | 50910 | 50909 |

[a] The secondary elements used were as follows: $A_1$, residues 5-23, 27-35, 39-53; $B_1$, residues 2-8, 13-19, 23-37, 43-46, 51-55; $B_4$, residues: 2-8, 13-19, 23-37, 42-46, 51-55.

[b] The parent structure for $A_1$ is $G_A$ (PDB 2FS1). The parent structure for $B_1$ and $B_4$ is $G_B$ (PDB 1PGA). RMSDs were calculated by superimposing the mean structure from the NMR ensemble with either the mean structure (in the case of 2FS1) or the X-ray structure (in the case of 1PGA) for the parent.

**Table S2**: Structure statistics for $S_{a1}$, $S_{b1}$, $S_{b2}$, and $S_{b3}$.

| | $S_{a1}$ | $S_{b1}$ | $S_{b2}$ | $S_{b3}$ |
|---|---|---|---|---|
| *A. Experimental restraint inputs* | | | | |
| NOE restraints | | | | |
| Sequential (i-j=1) | 89 | - | 35 | 40 |
| Medium range (1< i-j≤ 5) | 35 | - | 31 | 10 |
| Long range (i-j > 5) | 92 | - | 67 | 66 |
| Hydrogen bond restraints | 88 | - | 82 | 83 |
| TALOS dihedral angle restraints | - | - | - | 91 |
| Total NOE restraint inputs | 304 | | 215 | 290 |
| PRE restraints | | 41 | | |
| *B. Experimental chemical shift inputs* | | | | |
| $^{13}C\alpha$ | 88 | 79 | 83 | - |
| $^{13}C\beta$ | 86 | 70 | 76 | - |
| $^{13}CO$ | 69 | 67 | 72 | - |
| $^{15}N$ | 88 | 69 | 76 | - |
| $^{1}H_N$ | 88 | 69 | 76 | - |
| $^{1}H\alpha$ | 76 | 45 | 61 | - |
| *C. RMSDs to the mean structure (Å)* | | | | |
| Over all residues[a] | | | | |
| Backbone atoms | 2.77±0.82 | 5.47±1.86 | 2.34±0.60 | 2.46±0.64 |
| Heavy atoms | 3.51±0.85 | 6.32±1.86 | 3.00±0.62 | 3.29±0.61 |
| Secondary structures[b] | | | | |
| Backbone atoms | 1.07±0.23 | 3.79±1.50 (0.71±0.23)[c] | 1.08±0.24 | 0.68±0.14 |
| Heavy atoms | 1.84±0.37 | 4.37±1.42 (1.24±0.30)[c] | 1.78±0.32 | 1.42±0.25 |
| *D. Measures of structure quality (%)* | | | | |
| Ramachandran distribution | | | | |
| Most favored | 86.46±4.07 | 92.35±2.38 | 92.19±1.83 | 86.30±2.38 |
| Additionally allowed | 13.54±4.07 | 7.54±2.60 | 7.81±1.83 | 10.52±2.68 |
| Generously allowed | 0.00±0.00 | 0.00±0.00 | 0.00±0.00 | 1.47±1.54 |
| Disallowed | 0.00±0.00 | 0.12±0.38 | 0.00±0.00 | 1.96±1.10 |
| *E. Backbone RMSDs to the parent structure (Å)*[d] | | | | |
| Over all residues | 2.37 | 0.49 | 6.16 | 11.67 |
| Secondary structures | 0.88 | 0.41 | 3.39 | 3.20 |
| *F. PDB/BMRB codes* | | | | |
| PDB | 7MN1 | 7MQ4 | 7MN2 | 7MP7 |
| BMRB | 30901 | 30905 | 30902 | 30904 |

[a] Over all residues used as follows: $S_{a1}$, 1-95; $S_{b1}$, 4-85; $S_{b2}$, 1-93; $S_{b3}$, 1-87.

[b] The secondary elements used are as follows: $S_{a1}$, residues 2-10, 16-32, 40-44, 59-67, 72-81, 86-92; $S_{b1}$, residues 5-12, 17-24, 27-41, 46-50, 55-59, 73-83; $S_{b2}$, residues 2-9, 23-32, 43-48, 59-65, 71-80, 85-91; $S_{b3}$, residues 4-10, 24-37, 40-46, 51-57, 62-71, 79-85.

[c] RMSDs for $S_{b1}$ minus the putative $\alpha$2 region: residues 5-12, 17-24, 27-41, 46-50, 55-59.

[d] The parent structure for $S_{a1}$, $S_{b2}$, and $S_{b3}$ is PDB 1RIS. The parent structure for $S_{b1}$ is PDB 1PGA. In this case, the structure alignment is over the 56 amino acid B-region of $S_{b1}$.

**Supplemental Information**

**Comparison of $B_1$ and $S_{b2}$.** The sequences of the $B_1$ and $S_{b2}$ designs were based on aligning their β1-strands in the parent folds, and the corresponding ordered β1-regions of $B_1$ and $S_{b2}$ match reasonably well. In $B_1$, the β1-strand consists of amino acids 1-9. The equivalent residues in $S_{b2}$, amino acids 3-11, form most of β1 and part of the loop between β1 and α1. Beyond this region, however, there is substantial structural divergence for the corresponding residues in $B_1$ and $S_{b2}$ even though the sequence identity level is 80%. These topology differences are summarized here and in **Fig. S7A**. Amino acids 10-20 in $B_1$ form the β1-β2 loop and β2-strand, but in $S_{b2}$ the equivalent amino acids (residues 12-22) form a disordered loop. Amino acids 21-39 in $B_1$ form a 2-residue β2-α1 loop followed by a 15-residue α1-helix (residues 23-37) and part of the α1-β3 loop. By contrast, the corresponding amino acid sequence in $S_{b2}$ contains a 10-residue helix (residues 23-32) and nine residues of the α1-β2 loop. In $B_1$, amino acids 40-48 contain the 5-residue β3-strand whereas in $S_{b2}$ the equivalent region has a 6-residue β2-strand. Finally, amino acids 49-56 comprise the β4-strand in $B_1$ but form most of the disordered β2-β3 loop (residues 51-58) in $S_{b2}$. The 56 amino acid B-structure therefore undergoes significant topological changes when its sequence is embedded in the 93 amino acid S-type polypeptide chain.

Since the β1-strand contributes significantly to the cores of both $S_{b2}$ and $B_1$ and was used for aligning the sequences at the design stage, we analyzed how the contacts to β1 change between the two topologies (**Fig. S7B**). The β1-strand of $B_1$ is one of the central β-strands in the sheet and contributes the side chains of three amino acids, F3, L5, and L7, to the core. Other amino acids contacting these β1-strand core residues in the $B_1$ structure are T16, T18, and A20 (β2); A23, A26, F30, Y33, A34, and V37 (α1); V39 (α1-β3 loop); W43 and Y45 (β3); and F52 and V54 (β4). The corresponding core amino acids in the β1-strand of $S_{b2}$ are V5, I7, and L9. These residues make a nearly completely different set of contacts to those seen in $B_1$, having close interactions with Y58 (β2-β3 loop); L60, F62, and V64 (β3); L74 and L78 (α2); V84 (α2-β4 loop); and V87 and V89 (β4). Thus, the β1-core contacting residues in $S_{b2}$ are mostly outside the 56 amino acid region encoding the $B_1$-fold. Therefore, when the topologically aligned region is excluded, the cores of $B_1$ and $S_{b2}$ are largely non-overlapping, just as in the case of $A_1$ and $S_{a1}$. In analogy with the $A_1$/$S_{a1}$ system, the C-terminal end of the polypeptide chain beyond the B-coding region (residues 59-93) plays an important role in driving the folding process toward an S-fold.

**Rosetta calculations.** Rosetta energies of all designed structures were generated using the Slow Relax routine (2). 1000 decoys were calculated for each design. PDB coordinates and energy parameters for the lowest energy decoy for each design are included as supplemental files.

**Cloning.** Synthetic genes (Genescript) were cloned into a modified version of the pPal8 vector (Bio-Rad). The modified vector (denoted pA-YRGL) replaces the DNA sequence encoding the eXact tag with a DNA sequence encoding:

MEEAVDANSLAQAKEAAIKELKQYGIGDKYIKLINNAKTVEGVESLKNEILKALPTEGSGEEDKQYRGL-

**Protein Expression.** Cell growth is carried out either by IPTG methods or by auto-induction (3, 4). Cells were harvested by centrifugation at 3,750 x g for 20 minutes and lysed by sonication on ice in 0.1M $KPO_4$, pH 7.2. Cellular debris were pelleted by centrifugation at 10,000 x g for 15 minutes. Supernatant was clarified by centrifugation at 50,000 x g for one hour. High speed centrifugation is not essential, but will prevent columns from fouling with lipids and prolongs their life.

**Purification on the imidazole protease (*Im-Prot)* column (Potomac Affinity Proteins).** Loading and washing are at 5 mL/min for a 5mL *Im-Prot* column using a running butter of 0.1M $KPO_4$, pH 7.2. The amount of washing required for high purity depends on stickiness of the target protein and how much of it is bound to the column. We typically wash with 10 column volumes (CV) of wash solution followed by 3 CV high salt wash (e.g. 0.5M NaCl) and then ~ 10 CV running butter. This can be repeated as necessary. On an FPLC instrument, a very high uv absorbance is observed during the loading. During the washing with binding buffer, the uv absorbance will return to baseline. Injecting a shot of the high salt solution will release more uv-absorbing material from the column, followed by a return to baseline. The high-salt shots are repeated until the amount of absorbance release with each high-salt shot becomes small and constant. Make sure all the high-salt solution is washed out before initiating the cleavage. Different binding and washing buffers can be use as is appropriate for the target protein. The binding reaction itself is very strong and can be carried out under a variety of pH and salts.

**Triggering cleavage with imidazole.** The target protein is eluted from the *Im-Prot* column by injecting 15 mL of imidazole solution (0.1 mM) at 1mL/min). The cleaved protein should elute as a sharp peak in 2-3 CV. Make sure the column is at room temperature or the reaction will be considerably slower.

**Stripping the tag after elution.** Strip the tag by injecting 15 mL of 0.1 N $H_3PO_4$ (0.227 mL concentrated phosphoric acid (85%) per 100ml) at a flow rate of ~ 1 CV/min. Neutralize with wash solution immediately after stripping. DO NOT LET THE COLUMN SIT IN THE ACID for more than a few minutes.

**Storage.** Store columns at 4˚C in 20% Ethanol, 0.1M $KPO_4$, pH 7.2, 0.1mM EDTA. Do <u>not</u> store without buffer salts. The column should last at least a year if properly stored.

1. Y. He *et al.*, Structure, dynamics, and stability variation in bacterial albumin binding modules: implications for species specificity. *Biochemistry* **45**, 10102-10109 (2006).
2. A. Leaver-Fay *et al.*, ROSETTA3: an object-oriented software suite for the simulation and design of macromolecules. *Methods Enzymol* **487**, 545-574 (2011).
3. F. W. Studier, Protein production by auto-induction in high density shaking cultures. *Protein Expr Purif* **41**, 207-234 (2005).
4. P. A. Alexander, Y. He, Y. Chen, J. Orban, P. N. Bryan, A minimal sequence code for switching protein structure and function. *Proc Natl Acad Sci U S A* **106**, 21149-21154 (2009).
